# Transposable Element Expression and Sub-cellular Dynamics During hPSC Differentiation to Endoderm, Mesoderm, and Ectoderm Lineages

**DOI:** 10.1101/2024.07.03.602001

**Authors:** Isaac A. Babarinde, Xiuling Fu, Gang Ma, Yuhao Li, Katerina Oleynikova, Mobolaji T. Akinwole, Xuemeng Zhou, Alexey Ruzov, Andrew P. Hutchins

## Abstract

Transposable elements (TEs) are genomic elements that are found in multiple copies in mammalian genomes. TEs were previously thought to have little functional relevance but recent studies have reported TE roles in multiple biological processes, particularly in embryonic development. To investigate the expression dynamics of TEs during human early development, we used long-read sequence data generated from *in vitro* differentiation of human pluripotent stem cells (hPSCs) to endoderm, mesoderm, and ectoderm lineages to construct lineage-specific transcriptome assemblies and accurately place TE sequences in their transcript context. Our analysis revealed that specific TE types, such as LINEs and LTRs, exhibit distinct expression patterns across different lineages. Notably, an expression outburst was observed in the ectoderm lineage, with multiple TE types showing dynamic expression trajectories. Additionally, certain LTRs, including HERVH and LTR7Y, were highly expressed in hPSCs and endodermal cells, but these HERVH and LTR7Y sequences originated from completely different transcripts. Interestingly, TE-containing transcripts exhibit distinct levels of transcript stability and subcellular localization across different lineages. Moreover, we showed a consistent trend of increased chromatin association of TE-containing transcripts in germ lineage cells compared to hPSCs. This study suggests that TEs contribute to human embryonic development through dynamic chromatin interactions.

**Key findings:**

- Different loci of the same TEs are independently regulated in different cell states
- Ectoderm has the highest frequency of TE-containing transcripts
- The presence of TEs dynamically drives transcripts to different sub-cellular compartments in different cell states
- hPSCs have the least stable TE transcripts with the weakest TE chromatin association, highlighting loose hPSC chromatin and potential roles in cell differentiation

## Main

TEs are genomic elements with multiple copies resulting from autonomous and non-autonomous duplication in genomes ^1,2^. About half of the human genome consists of TEs of different types and properties^1,3,4^. TEs have multiple evolutionary histories with different genomic distributions across multiple mammalian species^5^. Because of their repetitive nature and their underrepresentation in protein-coding genes, TEs were previously considered to have little functional importance^6,7^. In many comparative genomics studies, TE-containing genomic regions were left out and considered as genomic dark matter^8,9^. However, recent studies have shown that TEs are functionally important in diverse biological processes^2,4^, including transcription^10–12^, post-transcription expression regulation^13,14^, transcript processing and stability^13,15,16^, chromatin regulation^17,18^, development^19,20^ and disease progression^21,22^. While TEs have been co-opted in normal developmental processes^23,24^, TEs still pose a risk to genomic integrity^4,25^. TEs are sources of mutations, can lead to genome rearrangement, interfere with normal regulatory networks and disrupt transcript coding potentials^4,26^. Therefore, TEs have both positive and negative aspects^7^. Although TEs have been studied in pluripotent stem cells^24,27^ and some terminally differentiated somatic tissues^21,28^, their expression patterns and their roles in post-implantation human development and gastrulation have not been well explored.

The presence of multiple copies of TEs in the genome makes the investigation of their functions difficult^29,30^. This is especially the case in TE expression quantification based on short-read RNA-seq in which a TE fragment often cannot be uniquely mapped to a genomic location^31^. Thus, assembling transcripts from only short-reads is fraught with difficulty even in well-annotated species such as human^32,33^, and attempting to include assembly of TE-containing transcripts is doubly difficult^31^. Previous efforts have considered global TE expression without consideration for the specific TE loci^21,34,35^, nor with their transcript context. Efforts are now being made to investigate TE expression at specific loci^27,29^, with an emphasis on understanding how TE sequences are spliced into transcripts. Locus-level expression quantification has been reported to be improved by transcript assembly^36^. Although both short-read and long-read can be used for transcript assembly^27,30^, the quality of the transcript assembly based on long-read data is superior^27,37^. Therefore, transcript assembly based on long-read data would substantially benefit TE expression studies.

We have previously shown that TE sequences are richly expressed in hPSCs, including inside coding and noncoding transcripts, and their presence is associated with changes in transcript biophysical processes^13^. It is generally assumed that TEs are most active in the pluripotent stage of development and decline in somatic tissues. However, this has not been explored, particularly in the context of TE sequences inside transcripts. In this study, we investigated human early development transcripts (hEDTs) assembled exclusively from long-read data to investigate TE expression dynamics in human early development (hED). We firstly differentiated hPSCs *in vitro* to the three germ layers; endoderm, mesoderm, and ectoderm, to mimic human gastrulation and conducted long-read RNA sequencing for each cell type. By integrating deepCAGE and polyA data, we showed that our transcripts are virtually complete from the 5’ to the 3’ ends of the transcripts. Using the assemblies, we showed that TE-containing transcripts were dynamically regulated across different loci and cell states. We next investigated the expression dynamics, subcellular localization, and transcript stability of the assembled hEDTs with a focus on the TE-containing transcripts and found that the stability and chromatin interaction of TE-containing transcripts increases during the differentiation process. Overall, this study presents the expression dynamics of TEs in early development and implicates TEs in cell state changes.

### hPSC transcriptome assembled from long-read RNA-Seq data

We have previously described an hPSC-specific transcriptome based on a combined long-read and short-read pipeline^27^. However, here we relied exclusively on deeply sequenced long-read data to assemble transcripts. This strategy has the advantage that TEs will be placed into their exact transcript context based only on long-reads, which will improve the accuracy of TE annotation in transcripts. To this end, we first generated PacBio long-read RNA-sequence data for hPSCs. Consensus reads were generated from the PacBio subreads, and then processed to produce noiseless reads using a standard PacBio read preparation pipeline (see Supplementary Methods) (https://github.com/nf-core/isoseq). The noiseless reads were then mapped to the human genome using Minimap2^38^. Next, StringTie2^39,40^ was used to assemble transcripts. After quality assessment, 33,375 hPSC transcripts, corresponding to 16,814 genes, were assembled.

Reference-based transcript assembly with long-read data are not completely error-free^32,41,42^. Errors might arise because of the long-read region-biased error^43^, reference-induced error^44^, or random RNA shearing during extraction. We, therefore, checked if our assembled transcripts were full-length transcripts supported by long-read data, as opposed to annotation-aided assembly. Indeed, the assembled transcripts were covered entirely by multiple consensus reads (**Fig. 1a**), suggesting that the transcripts were expressed in hPSCs. One advantage of transcript assembly based on long-read data only is the ability to sequence full-length mostly intact transcripts^30,31^. In support of this, most of the assembled transcripts were completely covered by at least one consensus read (**Fig. 1b**), suggesting that full-length transcripts were assembled. Interestingly, almost all the consensus reads used for the assembly were used without trimming (**Extended Data Fig. 1a**). As a negative control, we generated a ‘random GTF set’ with the same number and structure as the assembled transcripts (see Supplementary Methods). This random GTF was not supported by our long-read data (**Extended Data Fig. 1b**). The pileup of consensus reads shows that the reads map from the transcription start sites (TSSs) to the transcription end sites (TESs) of the hPSC assembly (**Fig. 1c**). The pileup of short-read data equally validates end-to-end transcript coverage for the hPSC assembly, but not for random GTF coordinates (**Extended Data Fig. 1c)**. These data highlight the power of long-reads in complete transcript assembly.

**Fig. 1:**
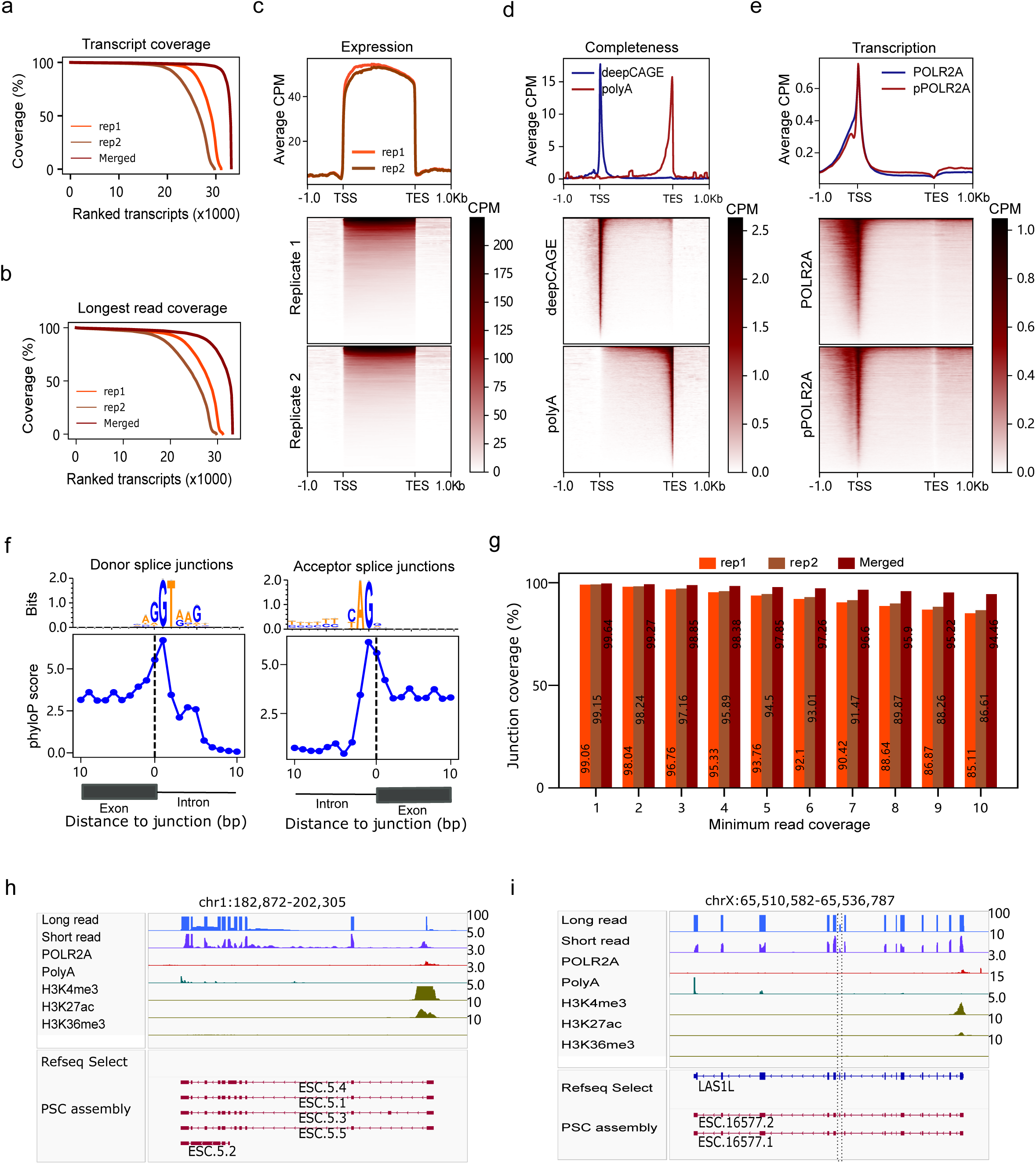
Quality assessment of the assembled hPSC long-read transcriptome. **a**, Consensus read coverage of the assembled PSC transcripts using all consensus reads. **b**, The longest consensus read coverage for the assembled PSC transcripts. **c**, Pileups for the consensus long reads for the assembled hPSC transcripts across the exons from the TSS to the TES. Each transcript is scaled to a uniform size. **d**, DeepCAGE (5’ end) and polyA tail (3’ end) pileups of the assembled transcripts, from the TSS to the TES scaled to a uniform size and not including intronic regions. PolyA data were from GSE138759 and GSE111134 while deepCAGE data were from GSE34448 and GSE61264. **e**, Density pileup of POLR2A and phosphorylated POLR2A (pPOLR2A) for the assembled hPSC transcripts. The data were from GSE242645. **f**, Position weight matrix nucleotide frequencies and phyloP conservation scores of the 10-bp positions around the donor and acceptor splice junctions of the assembled multi-exon hPSC transcripts **g**, Proportion of hPSC junctions covered by indicated minimum number of reads. Integrative genomics views showing multiple novel transcripts in a locus (**h**) and a variant transcript made by a skipped exon of LASIL gene (**i**). The position of a skipped exon is marked by dotted line in **Fig. 1i**.

To further assess the completeness of the assembled transcripts, we utilized published hPSC deepCAGE^45^ and polyadenylated (polyA) data that marks the TSS and TES, respectively. The long-read assembled TSSs were enriched for deepCAGE signal tags, while the TESs were enriched for the polyA signal (**Fig. 1d**). These enrichments were not found in the random GTF set (**Extended Data Fig. 1d)**. Similarly, ChIP-Seq data of hPSC POL2R, H3K27ac and H3K4me3 showed the expected promoter enrichment in the hPSC assembly, but not in the random GTF (**Fig. 1e, Extended Data Figs. 1e, f)**. Additionally, H3k36me3 signal is enriched on the hPSC transcript bodies (**Extended Data Fig. 1g**). The specific RNA-seq and histone modification data extensively validate the completeness of the hPSC assembly.

We next checked if the splice junctions of the assembled transcripts were supported. The evolutionary conservation scores across the junctions were higher relative to introns. The highest conservation was in the dinucleotide at the splice junctions for both the 5’ and 3’ ends (**Fig. 1f**). Indeed, the nucleotide frequencies showed the typical GT dinucleotide at the donor site, and AG at the acceptor site. Analysis of short-read RNA-seq data with an anchor length of 10 bp, showed that more than 99% of the splice junctions have at least one short-read junction support, and 85% have 10 or more reads (**Fig. 1g, Extended Data Fig. 1h)**. Indeed, out assembly correctly identified transcripts of *POU5F1* and *SLC2A* with the necessary deepCAGE, POLR2A, polyA and epigenetic peaks (**Extended Data Figs. 1i, j**). Additionally, our assembly included previously unannotated novel transcripts such as a cluster of transcripts found on human chromosome 1 (**Fig. 1h**). We were also able to assemble new variants of known genes, such as an isoform of *LASIL* gene with a skipped exon (**Fig. 1i**). Taken together, these results, from multiple data, demonstrate the fidelity of the terminals and the splice junctions of the assembled transcripts.

### *In vitro* differentiation drives consistent and specific transcriptome trajectory in germ layers

To investigate transcript expression dynamics during human embryonic development, we performed hPSC *in vitro* differentiation to three germ layers using previously published protocols^46^ (**Fig. 2a**). Lineage-specific marker genes were specifically up-regulated in each lineage while pluripotent markers gradually decrease along the differentiation time course (**Extended Data Fig. 2)**. To further confirm the fidelity of our differentiation experiments, we performed Fluorescence-Activated Cell Sorting (FACS) using *PAX6*, the marker for ectoderm cells. As expected, the success of the ectoderm differentiation was confirmed by the expression of *PAX6* (**Fig. 2b**). Compared to 1% in PSC, >90% of the ectoderm cells expressed *PAX6*. After the initial confirmation of our cell types, long-read sequencing was done for two technical replicates for each cell state, with each sample sequenced to 20-45 million PacBio raw subreads (**Extended Data Fig. 3a)**. The consensus reads generated from each replicate ranged from 509 thousand to 1.2 million. Individual assembly produced 31,245 endoderm, 37,161 mesoderm, and 56,063 ectoderm transcripts (**Fig. 2c**). These corresponded to 14,989 endoderm, 17,520 mesoderm, and 24,766 ectoderm genes (**Extended Data Fig. 3b**). The saturation analyses with down-sampling suggest that increasing the sequencing depths would lead to the discovery of more transcripts, especially the novel transcripts (**Extended Data Fig. 3c**), consistent with the previous report^47^. Since the number of assembled transcripts would be affected by the sequencing depth^31^, we asked if the higher transcript count in ectoderm cells is a genuine phenomenon or a consequence of greater sequencing depth in these samples. Interestingly, transcript assembly with individual ectoderm samples produced more transcripts even though the sequencing depths were lower than the combined sequencing depths in other cell states (**Extended Data Figs. 3a-b)**, suggesting that transcript complexity is higher in ectoderm cells. To further confirm this, we subsampled 300,000 consensus reads from the replicates of each cell state. Transcript assembly from the uniform read counts revealed that more transcripts were assembled in ectoderm (**Extended Data Figs. 3d**). As the impact of sequencing depth tends to be more pronounced in lowly expressed transcripts^31^, we checked if the higher transcript count in ectoderm would persist at a higher expression threshold. Indeed, that was the case up to 20 counts per million (CPM) expression threshold (**Extended Data Figs. 3e**), supporting the increased complexity of ectoderm transcripts.

**Fig. 2:**
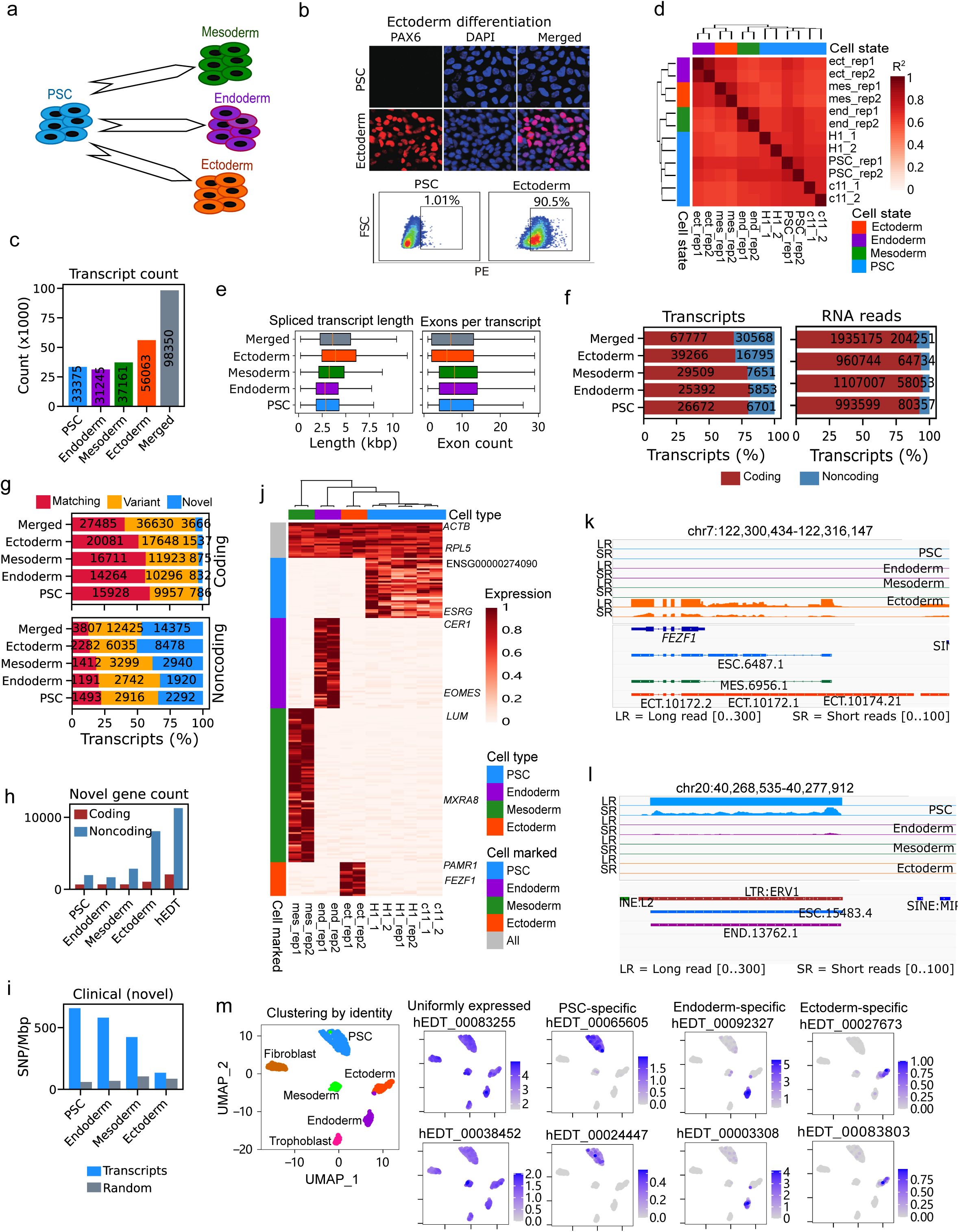
Expression dynamics of the assembled human early development transcripts. **a**, Simplified experimental design for the *in vitro* differentiation of human hPSC to each of the three embryonic germ layers. The detailed experimental procedures are in the Supplementary Methods. **b**, Fluorescence-activated cell sorting results using PAX6 to demonstrate the successful differentiation of PSC to ectoderm. **c**, The numbers of transcripts assembled from each cell state and the merged transcript assembly. **d**, The expression correlation heatmap of the samples made from long-read quantification. H1_1, H1_2, c11_1 and c11_2 samples were from previously reported data^27^. **e**, Boxplots showing the spliced transcript lengths (left) and number of exons per assembled transcript (right) for the assembled transcripts. Boxplots show the mean (red central bar), and second and third quartiles, and the whiskers show 1.5 times the interquartile ranges. Kruskal-Wallis p value = 2.37 × 10^-28^for spliced transcript lengths and Kruskal-Wallis p value = 1.86 × 10^-173^ exon per transcript. **f**, Stacked bar chart showing the proportions and numbers of coding and noncoding transcripts in the hEDT assemblies. The left plot shows the distribution of the assembled transcript counts while the right plot shows the distribution of the total RNA read counts from the long-read quantifications. **g**, Stacked bar chart showing the proportions and numbers of assembled transcripts based on the similarity to version 43 of the GENCODE assembly. Matching transcript completely matches to a known transcript (including all exons and splice junctions), while a variant transcript partially overlaps a known GENCODE transcript. Novel transcript does not overlap any known GENCODE transcript exon. **h**, The number of novel coding and noncoding genes identified in each cell state. Because a gene might have more than one transcript, a gene could both be coding and noncoding. **i**, The enrichment of SNPs with clinical relevance in novel transcripts. The enrichment for matched background sequences (with equal lengths and exon structures are also shown). In all cell states, novel transcripts had significantly higher enrichment of clinical SNPs (Fisher Exact p value < 0.001). **j**, Heatmap showing the uniformly expressed transcripts and transcripts specific to each cell state based on the long-read quantification. For each transcript, the expression was normalized by the highest expression level. Integrative genomics views showing the expression of ectoderm enriched transcripts (**k**) and PSC-enriched novel transcript (**l**). **m**, Single-cell RNA-seq expression showing uniformly expressed transcripts and marker transcripts specific to the indicated cell states.

Transcript quantification using the merged transcript reference showed that samples from the same cell state tended to cluster together and separated from other samples (**Fig. 2d, Extended Data Fig. 3f)**. Importantly, the differentiated samples were all separated from hPSC samples, and did not cluster with samples from other cell states, suggesting that different cell states had a specific transcriptomic landscape. Indeed, previously generated long-read data of H1 and c11 cell lines^27^ consistently clustered with hPSC data, demonstrating the robustness of our pipelines to technical replicates and different hPSC cell lines. To further confirm the fidelity of our differentiation experiments, we retrieved 16 human iPSC, endoderm, mesoderm and ectoderm RNA-seq data sequenced on Nanopore platform^48^. We performed transcript quantification based on our merged hEDT assembly, and performed pairwise correlations between our samples and the Nanopore samples. The correlation analyses showed that the closest Nanopore sample had the same identify for all the eight PacBio samples (**Extended Data Fig. 3g)**. These results demonstrated the reliability of our *in vitro* differentiation experiments and transcript assembly and quantification procedures.

We then proceeded to investigate the features of hEDT across different cell types. Ectoderm transcripts tended to have longer spliced transcripts with fewer exons per transcript (Mann-Whitney U p value < 1 × 10^-5^) (**Figs. 2 e**). However other parameters, including un-spliced transcript lengths, exon lengths and numbers of transcripts per gene were not substantially different across different cell states (**Extended Data Fig. 3h)**. We next predicted coding and noncoding potential using FEELnc^49^, and transcript distributions based on protein coding potential showed that ectoderm tended to have more noncoding transcripts and expressed more noncoding RNA reads (**Fig. 2f**). The comparisons of the assemblies of each cell states to the GENCODE (v43) transcript set^50^ revealed that ectoderm indeed had a lower proportion of GENCODE-known (matching) transcripts (**Extended Data Fig. 3i)**. Noncoding transcripts tended to be novel transcripts in all cell types, with a higher proportion of novel transcripts in ectoderm (**Fig. 2g**). Even at gene level, the majority of the novel genes were noncoding (**Fig. 2h**). These results highlighted the ability of assembly to identify previously unknown transcripts.

With thousands of novel transcripts, we checked for the signatures of functionality for the novel transcripts. Compared to the random sequences, both novel and matching transcripts were found to be enriched in SNPs with clinical and functional relevance (**Fig. 2g, Extended Data Fig. 3j**), however higher enrichment was found for matching transcripts. Although ectoderm enrichment was lower, the novel transcripts from all cell states had significantly higher functional SNP enrichment compared to matched random sequences (Fisher Exact p value < 0.001). We then checked if novel and matching transcripts had similar transcription factor binding motif (TFBM) enrichment in the core promoters of the transcripts. Indeed, the correlations of the TFBM enrichments between novel and matching transcripts were >0.6 for the four cell states (**Extended Data Fig. 3k**). Although the majority of the significant transcription factors (TFs) were found in both matching and novel transcripts, some TFs were found to be enriched only in novel or matching transcripts (**Extended Data Fig. 3l**), highlighting both the presence of both unique and shared transcription factors for matching and novel transcripts. These data suggest that a substantial subset of the assembled novel transcripts was functionally relevant.

To further check the reliability of the assembled transcripts, we checked the splice sites of the assembled transcripts. Across cell states, base-level conservation levels and nucleotide frequencies of the splice junctions showed that the assembled transcripts were reliable (**Extended Data Figs. 4a, b)**. The distributions of the alternative splicing events were found to be similar across the four cell states (**Extended Data Fig. 4c**). The consistency of the splicing signals and the similarity across cell states suggest that the quality of our assembled transcripts was reliable.

To investigate the overall expression dynamics of the assembled transcripts, we extracted lineage specific markers and uniformly expressed genes across the four cell states (**Fig. 2j**). The uniformly expressed genes included known housekeeping genes such as *ACTB* and *RPL5,* while the expressions of the known lineage specific markers were confirmed. As an example, we confirmed that ectoderm-specific *FEZF1* had both long-read and short-read supports for ectoderm specificity (**Fig. 2k)**. *EOMES*, a known endoderm marker was found to be expressed only in the endoderm with both long and short read support (**Extended Data Fig. 4d**). A mesoderm marker, *IGFBP3*, was found to be exclusively expressed in the mesoderm (**Extended Data Fig. 4e**). Interestingly, we also found some novel transcripts such as ESC.15483.4 that was highly expressed in PSC with both long and short read supports (**Fig. 2l**). As expected, the unform expression of *ACTB* was supported with both long and short reads. The different expression patterns could be as a results of different cell states or the consequences of the differentiation factors used in the experiments. To further investigate this possibility, we subjected the human embryonic skin fibroblast (hESF) cells to mesoderm and endoderm differentiation conditions. Unlike in PSC, all hESF-derived samples clustered together, suggesting that hESFs did not respond to the differentiation process (**Extended Data Fig. 4e**). This result showed that the expression dynamics of the cells actually reflected the cell states, and not just a reflection of medium treatment.

To investigate the heterogeneity of transcript expression at the single cell level, previously published scRNA-seq data^51^ containing hPSCs, and differentiated cell types was reanalyzed against our hEDT assembly. UMAP clustering revealed that cells of similar states tended to cluster together (**Extended Data Fig. 5a**). Meanwhile, hEDTs specifically expressed in each cell state and the top 20 uniformly expressed transcripts were extracted from the long-read data (**Fig. 2j**). We then checked the expression of the selected hEDTs and found that the expression patterns were consistently reflected in scRNA-seq cell states (**Fig. 2m, Extended Data Figs. 5b-f)**. Both bulk (long-read and short-read) and scRNA-seq data largely consistently showed that the *in vitro* differentiation and bioinformatics procedures reliably captured the expressed hEDT set.

### Different TE types are dynamically spliced into coding and noncoding transcripts during *in vitro* differentiation

Having established the reliability of the *in vitro* differentiation experiments and the assembled transcripts, we then investigated TE splicing patterns across the cell states. We used *nhmmer* to annotate TEs inside transcript sequences based on a previously described approach^2,52^. The proportions of TE-containing transcripts and RNA read in the assembled transcripts revealed that ectoderm expressed more TE-containing transcripts (**Extended Data Fig. 6a**). As previously reported^27^, most noncoding transcripts tended to contain more TEs than the coding transcripts (**Fig. 3a)**. This observation was true when we considered the number of TE-containing transcripts and total TE-containing RNA count, highlighting the coding potential disruption ability of TEs. Notably, in both coding and noncoding transcripts, ectoderm had more TEs. Intriguingly, most TE-containing transcripts contained less than half TE-derived sequence, and only a small proportion of the transcripts contained >50% TE-derived sequences (**Fig. 3b**). Again, there was a stark difference between coding and noncoding transcripts. TE-containing noncoding transcripts contained more TE-derived sequences, although most noncoding transcripts still contained less than 50% TE-derived sequences. The distributions of TE coverage were more similar in coding transcripts than in noncoding transcripts. Specifically, ectoderm TE-containing noncoding transcripts tended to contain more intermediate values. However, hPSC and endoderm contained the highest proportion of transcripts consisting entirely of TE-derived sequence (**Fig. 3b**) even though they had fewer overall TE-containing transcripts (**Fig. 3a**), highlighting the differences across cell states.

**Fig. 3:**
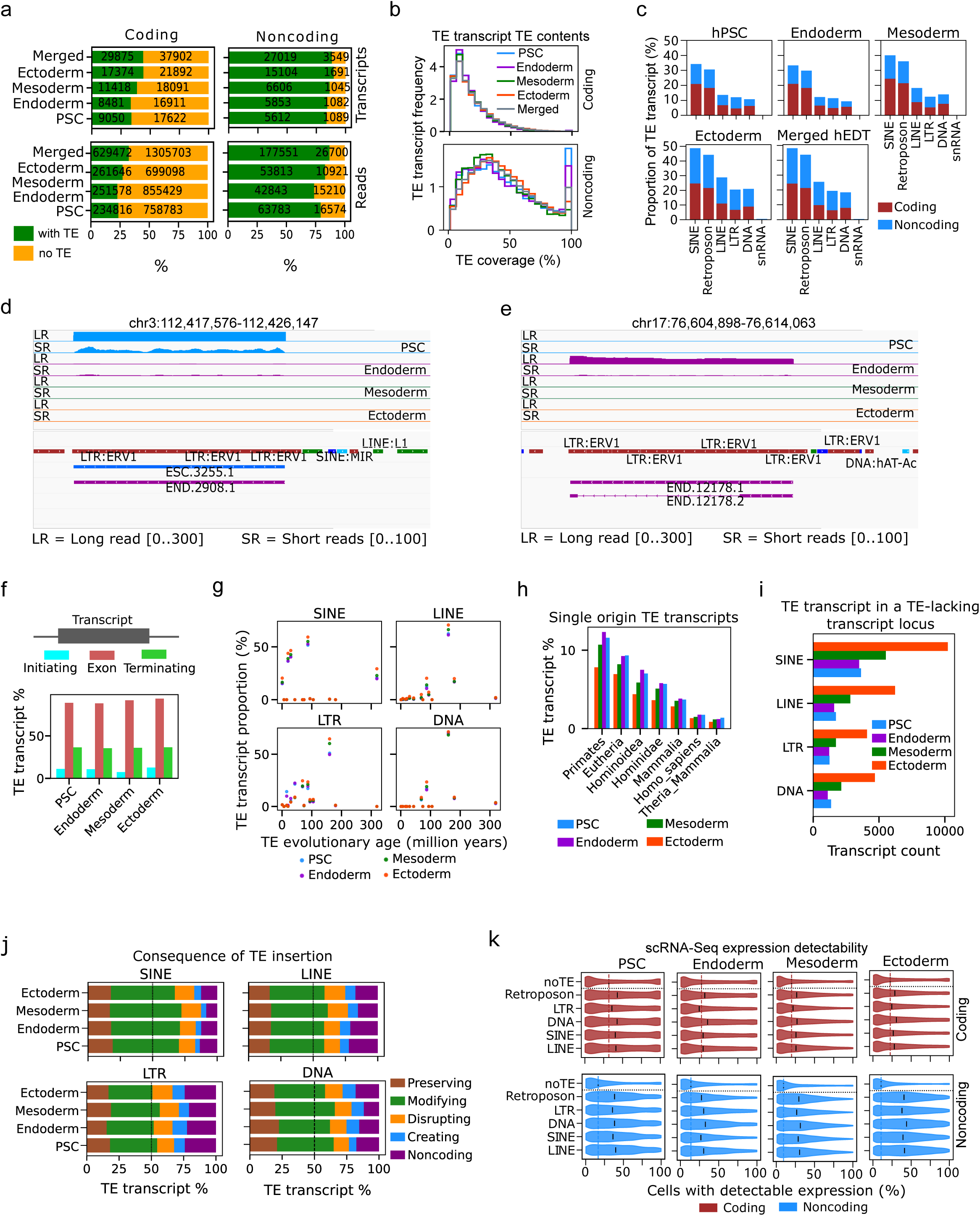
Divergent TE expression dynamics across coding and noncoding transcripts. **a**, Stacked bar charts showing the proportion and number of TE-containing transcripts in the coding (left) and noncoding (right) transcripts in the hEDT assemblies. The upper plots show the assembled transcript counts while the lower plots show the RNA read counts from long-read transcript quantification. **b**, Histograms showing the distributions of TE coverages (percent of sequence that consists of TE) for both coding and noncoding transcripts. **c**, Stacked bar charts showing the proportion of TE-containing transcripts for coding and noncoding transcripts in the assembled hEDTs. Integrative genomics views showing different loci of PSC-enriched (**d**) and endoderm-enriched (**e**) LTR-containing novel transcripts. **f**, Bar plots showing the proportion of TE-initiating, -exonized and -terminating transcripts. g, Scatter plots showing the proportion of TE transcripts that evolved at different evolutionary timepoints. The evolutionary ages were estimated based on the phylogeny of the species for the TE. **h**, Proportion of TE-transcripts of single age category. **i**, The number of TE-containing transcripts with at least one TE-lacking transcript of the same gene. **j**, The consequences of TE insertion based on the comparison between TE-containing and the closest TE-lacking transcript of the same gene. **k**, Single-cell RNA-seq expression detectability of different groups of transcripts within the cell states. The red and blue dotted lines represent the median values for the coding and noncoding transcripts without TEs.

To investigate the specific TE splicing pattern further, we checked which TE types were contained in hEDTs. We found that the most frequently spliced TE types included SINE, retroposon, LINE, LTR, and DNA (**Fig. 3c**). It is important to note that these TE frequencies did not reflect the genomic TE distribution (**Extended Data Fig. 6b**), suggesting the TEs are not just randomly spliced from genomic sequences. We found that the majority of SINE-containing transcripts also had retroposon motifs, suggesting that these two TE types co-exist in hEDTs (**Extended Data Fig. 6c**). Consequently, SINE-containing transcripts were considered to represent retroposon-containing transcripts in the clustering analyses. Across the major TE types, frequent TE splicing in ectoderm was observed with almost half of the transcripts containing at least one SINE, compared to about a third in hPSC transcripts (**Fig. 3c**). Indeed, the higher frequency of SINE and retroposon splicing is observed across cell states in both coding and noncoding transcripts. Interestingly, a number of TE-containing transcripts were expressed in a cell state-specific manner. For example, a locus containing a cluster of LTRs on human chromosome 3 is exclusively expressed in PSC (**Fig. 3d**) while another LTR-containing locus on human chromosome 17 was expressed exclusively in endoderm (**Fig. 3e**). Indeed, multiple cell state-specific TE-containing transcripts such as endoderm-specific (**Extended Data Fig. 6d**), ectoderm-specific (**Extended Data Fig. 6e**) and mesoderm-specific (**Extended Data Fig. 6f**) were detected, highlighting cell state-specific TE expression.

Since only a small proportion of transcripts are entirely covered by TEs (**Fig. 3b**), we asked which part of the transcript TE is located. In most of the transcripts, TEs are inserted into the transcript exon; a sizable number of transcripts were TE-terminating (having TE on their TTS) while very fewer transcripts have TE-overlapping TSS (**Fig. 3f**). Across different TE types, the prevalence of exonized TE transcripts were consistent but the proportions of TE-initiating and TE-terminating were heterogenous (**Extended Data Fig. 6g**). In many cases, we found that a transcript may contain more than one TE. This is especially the case for SINE-containing transcripts with more than 50% containing multiple SINEs (**Extended Data Fig. 6h**). However, higher proportion of DNA-containing transcripts contained single TE. The analyses of the evolutionary times of inserted TEs showed that TEs of different evolutionary ages tended to be found in the same transcript (**Extended Data Fig. 6i**). This is especially so for SINEs but less so for DNAs for which more than 80% of DNA-containing transcripts had single-origin TE. Interestingly, although different TE types inserted into the transcripts seemed to have different evolutionary times, the overall patterns were similar across cell states (**Fig. 3g**). Focusing on TEs with single evolutionary origin, we found that majority of the TEs evolved in the Primates common ancestor (**Fig. 3h**). The next most common evolutionary time was in the eutherian common ancestor. Interestingly, the uniqueness of the evolutionary history of SINE-containing transcripts became more obvious when single-origin TE transcripts were considered (**Extended Data Fig. 6j**). Specifically, unlike for SINEs, the majority of the expressed LINEs, LTRs and DNAs evolved in Eutheria common ancestor, revealing the heterogeneity of TE evolutionary histories.

To investigate the consequences of TE insertion into transcripts, we focused the analyses on loci containing both TE-lacking and TE-containing transcripts. Consistent with the overall pattern, ectoderm cells tended to contain more TE-containing transcripts with TE-lacking splice variants, and SINE-containing transcripts tended to be more (**Fig. 3i**). We then attributed TE-containing transcripts to the closest TE-lacking transcripts. Interestingly, more than 75% of the TE-containing transcripts were either coding or linked to a protein-coding TE-lacking transcripts (**Fig. 3j**). LTR-containing transcripts tended to have more TE-containing noncoding transcripts for which the closest TE-lacking transcript was also noncoding. Across TE types and cell states, the major consequence of TE insertion was “modifying” in which the TE insertion substantially affected the CDS length but did not lead to the loss of coding potential. Interestingly, we also found instances in which TE insertion led to the complete loss (disrupting) or gain (creating) of coding ability. These results suggest that TE insertion may have different consequences, especially on the coding potential, of the TE-lacking transcripts.

Because TE insertion could have serious consequences on the coding potentials, we investigated the expression dynamics of TE-containing coding and noncoding transcripts. For both coding (**Extended Data Fig. 7a**) and noncoding (**Extended Data Fig. 7b**) transcripts, SINEs and retroposons were more predominant. Whereas about 80% of noncoding transcripts in ectoderm contain a SINE element, less than 10% of the coding transcripts contained LTR-containing transcripts. The coding bias of the TEs was evident as the proportion of coding transcripts varied across cell states and TE types, ranging from 33% for LTR in ectoderm to 63% for SINE in endoderm (**Extended Data Fig. 7c**). While most of the TE-free transcripts were either known GENCODE transcripts (matching) or new isoforms of known GENCODE transcripts (variants), TE-containing transcripts, especially the noncoding transcripts, contained more novel transcripts (that do not overlap any known GENCODE transcript) with the highest novel proportion (∼60%) for DNA TE-containing transcripts in ectoderm noncoding transcripts (**Extended Data Fig. 7d**). As TEs could potentially create a minor transcript as an alternative isoform of a major functional transcript, we investigated if TE-containing transcripts tended to contain fewer major transcripts (transcripts with the highest expression among the isoforms of a single gene). For coding transcripts, the TE-free sequences tended to have a higher proportion of major transcripts, compared to the TE-containing transcripts (**Extended Data Fig. 7e**). Conversely, TE-containing noncoding transcripts tended to contain a higher proportion of major transcripts, suggesting that many TE-containing coding transcripts were the products of alternative splicing. The data demonstrated that our long-read assembly uncovered multiple previously unannotated transcripts and the bias of the insertion of various TE types into transcripts.

TEs might contribute to cell states or serve as functional modules for cell state transitions^53,54^. As previously reported^55^, noncoding transcripts had higher expression variability, as quantified by the coefficient of variation, across cell types (**Extended Data Fig. 7f**). Further, TE-containing coding and noncoding transcripts tended to have higher expression variability than TE-free transcripts. Interestingly, the Tau coefficient which measures expression specificity showed that both TE-containing coding and noncoding transcripts had higher cell-specific expression than TE-free transcripts, suggesting that TE expression is regulated in a cell type-specific manner. We next checked the expression variability within cell states using scRNA-seq data and found that hPSC transcripts had lower expression variability than differentiated cells (**Extended Data Fig. 7g**). Except for mesoderm LTR-containing transcripts, noncoding transcripts tended to have higher expression variability than the coding transcripts. Notably, TE presence led to higher expression variability in coding transcripts but lower variability in noncoding transcripts. The differences in the expression variability might be influenced by the expression detectability, especially for lowly expressed transcripts. We checked the expression detectability and found that TE-containing transcripts tended to have higher detectability (**Fig. 3j**). Interestingly, while the transcript detectability in TE-free noncoding transcripts was lower than that of the TE-free coding transcripts, the detectability of the TE-containing coding and noncoding transcripts were more similar. These data showed that the low expression variability in TE-containing noncoding transcripts was not just because of the inability to detect due to the low transcript expression. Taken together, these results suggest that the trajectory of TE expression is coordinated during differentiation, but cell-level coding and noncoding transcript expression is dynamic.

### Biased TE splicing patterns and frequencies of TE sequences in coding and noncoding transcripts

Having established the dynamic expression of TE-containing transcript in different lineages, we investigated the splicing patterns and insertion frequencies of different TEs in the four cell states. Because of the differences in coding and noncoding transcripts, we investigated the splicing patterns separately. Analysis of TE-splicing patterns in coding transcripts showed that TE sequences are rare in coding sequence (CDS) regions (**Fig. 4a, Extended Data Fig. 8a**), presumably reflecting the evolutionary cost of a TE inserting into and disrupting a coding sequence. Across TE types and cell states, TEs were overrepresented in the 5’ and 3’ untranslated regions (UTRs) compared to the CDS, with higher frequencies in the 3’ UTRs. However, investigation of specific TEs showed differences between different regions and cell states. For example, while HERVH and MSTB are both LTRs, their TE splicing patterns were different, with HERVH sequences being rare in the 3’UTR, whilst MSTBs were enriched (**Fig. 4a**). In addition, there was a higher frequency of HERVH TE-derived sequences in endoderm cells. Interestingly, the TE pattern of X24_LINE was very similar to that of MSTB, and also showed higher splicing in the UTRs of ectoderm, revealing the heterogeneity of the splicing patterns of specific TEs. The comparison of the MER3 and LTR6A frequencies further highlighted differences between these TEs and different cell states (**Extended Data Fig. 8a**).

**Fig. 4:**
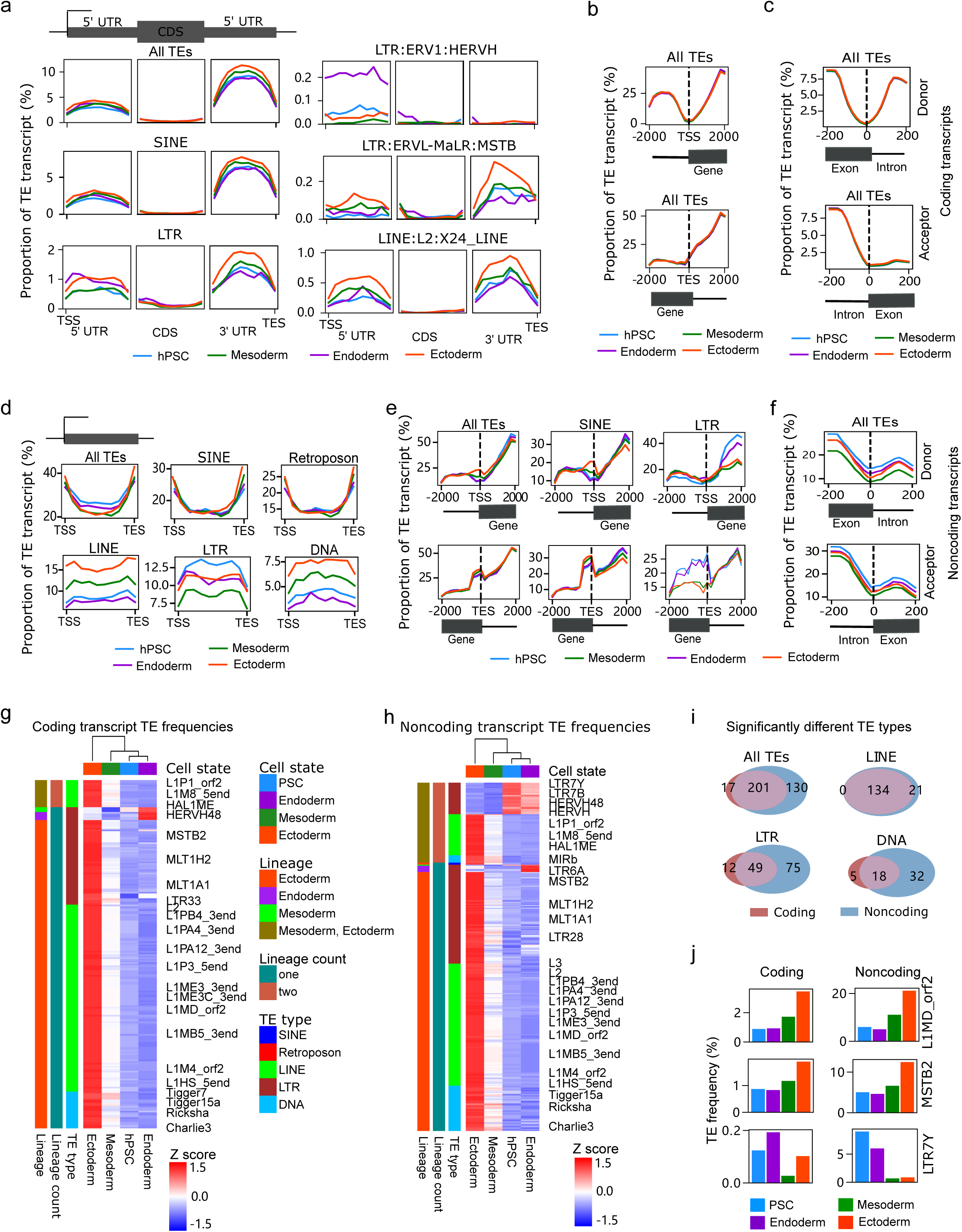
TE splicing patterns and frequencies of TE-containing coding and noncoding transcripts. **a**, The splicing patterns of different groups of TEs in coding transcripts. Coding transcripts were divided into 5’ UTR, CDS and 3’ UTR, and scaled to a uniform size. **b**, Proportion of TEs 2 kbp upstream and downstream of coding transcript TSS and TES. **c**, Proportion of TEs around 200 bp upstream and downstream of the donor and acceptor splice sites of coding transcripts. **d**, The splicing patterns of different TE types in noncoding transcripts. Each transcript is scaled to a uniform size. **e**, TE contents around the TSS and TES of noncoding transcripts. **f**, TE contents around the donor and acceptor sites in the noncoding transcripts. For panels **a-f**, each transcript was divided into 20 bins. The TE content for each bin was then computed. **g**, Heatmap of TEs with significant difference in coding transcript frequencies between hPSC and any of the three germ layers. **h**, Heatmap of TEs with significant frequency differences in the conceding transcripts of the hPSC and any of the three germ layers. For panels **g** and **h**, the statistical significance was defined as an adjusted p-value < 0.05 and odds ratio ≥ 2. For the heatmap, the z-score of the proportion of TE-containing coding and noncoding transcripts was computed. For each TE, the lineage(s) with significant difference, and the number of lineage(s) with significant differences are shown. **i**, Venn diagrams of the overlaps of the TE types with significant frequency differences in coding and noncoding transcripts. **j**, TE frequencies of selected TEs in the coding and noncoding transcripts.

We next looked at splicing at the TSS and TES boundaries of transcripts, and interestingly, while the TE splicing patterns across the TSS and TES were different, there were no substantial differences across cell states (**Fig. 4b**). Across the TE types, TE proportion was low at the TSS. In contrast, TEs were also reduced at the TES, although this reduction extended 5’ into the transcript body (**Extended Data Fig. 8b**). Similarly, TE frequencies at both the donor and acceptor splice junctions were low (**Fig. 4c, Extended Data Fig. 8c**). Although there are subtle differences across cell lines and TE types, the general patterns were similar (**Extended Data Fig. 8c**). Surprisingly, TE frequencies of the exons at the splice sites suggest that TE insertion suppression at the exon starts is stronger than the suppression at the exon ends. This supports the idea that TEs are impaired from splicing into TSSs, TESs, or splice junctions directly to prevent the disruption of important motifs for transcription initiation and termination and transcript splicing.

We next studied the properties of TEs in noncoding transcripts and found that TE splicing patterns in noncoding transcripts were dynamic between TE types, cell states, and even positions within the transcript (**Fig. 4d, Extended Data Fig. 8d**). Specifically, while SINE and retroposon TEs are enriched towards the 5’ and 3’ ends of the transcripts, LINE, LTR, and DNA TEs are uniformly distributed. Additionally, the overall frequencies were different across cell states (**Fig. 4d, Extended Data Fig. 8d**). Ectoderm noncoding transcripts were enriched for LINE and DNA TEs, whilst LTRs were enriched in hPSCs, and were also high in ectoderm and endoderm (**Fig. 4d**). The dynamics of TE expression was more obvious for transcripts containing LTRs. For example, while HERVFH21 was concentrated in the center of noncoding transcripts, endoderm LTR6A had higher enrichment around 5’ and 3’ ends, suggesting these transcripts are noncoding remnants of intact ERVs. HERVH, conversely, was uniformly distributed across the noncoding transcripts in hPSC and endoderm cells (**Extended Data Fig. 8d**), highlighting the differences among TEs.

TE frequencies around noncoding transcript TSSs were also dynamic. SINE, LTRs, and retroposon were rare at the TSS (**Fig. 4e, Extended Data Fig. 8e)**. Conversely at the transcript ends, SINEs and LTRs were enriched up to the TES, and then their enrichment dropped substantially, suggesting that they marked transcript ends. TE sequences were absent at the TSSs of noncoding transcripts, suggesting TE sequences rarely act as core promoters of TSSs. TE frequencies at noncoding transcript splice junctions were also low (**Fig. 4e, Extended Data Fig. 8f)**, but not as low as in coding transcripts (**Extended Data Fig. 8f)**. Across the noncoding transcript bodies, TSS, TES, and splice junctions (**Figs. 4d-f, Extended Data Figs. 8d-f**), splice pattern differences based on the TE type and cell states were obvious.

To figure out the cell type-specific TE activities, we explored their frequencies in coding and noncoding transcripts. The comparisons of the frequencies of TE-containing coding transcripts identified 218 TEs that were significantly different in their frequencies upon differentiation into endoderm, mesoderm, or ectoderm (**Fig. 4g**). The majority of the TEs that were significantly different were enriched in ectoderm cells and were LINEs and LTRs and a few DNA TEs. In contrast, some LTRs like LTR7, HERVH, HERVH48, and MTSB2 were enriched in endoderm coding transcripts. On the other hand, the frequency comparison for noncoding transcripts identified 331 specific TEs that were differentially enriched in at least one of the four cell states (**Fig. 4h**). Similar to the coding transcripts, the majority of the differentially spliced TEs in noncoding transcripts were LINEs and LTRs, and they were more frequent in the ectoderm. Some LTRs such as LTR7B and HERVH were more frequent in hPSCs as previously reported^56,57^ and here we observed many of them in endoderm cells, suggesting these are not wholly specific features of hPSCs. Indeed, LTR7B/Y/H and HERVH were enriched in endoderm cells (**Fig. 4h**), supporting the idea that HERVH expression extends to cell types beyond hPSCs. HERVH modulates 3-dimensional genome structure in hPSCs^58^ and may also be performing this function in somatic cells. Interestingly, there was a substantial overlap in the significantly enriched TE types in any cell type for both coding and noncoding transcripts (**Fig. 4i**). Breaking this down by TE type, the overlap is high for LINEs, with just 21-specific for noncoding transcripts and 134 LINE types were enriched in both coding and noncoding. For example, although the frequencies were different, the overall abundance patterns for LINE L1MD_orf2 were similar for coding and noncoding transcripts (**Fig. 4j**). In contrast to LINEs, LTRs were different between coding and noncoding transcripts, and only 49 LTR-types were common. This was exemplified by the LTRs

MSTB2 and LTR7Y which were specific to cell types **(Fig. 4j)**. Overall, these data indicate that the TE content in different lineages is highly dynamic, whilst the TE sequences in coding and noncoding transcripts are more similar, except for LTRs, which are cell type and transcript-type specific.

### TE-containing coding and noncoding transcripts expressed from different genomic loci are independently regulated

Above, we mainly consider TEs at the family or type level. However, a significant advantage of long-read sequence data is that we can place TEs into their specific transcript context. Although there is a substantial overlap in the types of TEs that are found in coding and noncoding transcripts, the exact genomic loci or the TE-containing transcripts are likely to be different. Potentially there are two scenarios of TE expression in different cell states. The first scenario is that the same TE type is contained in the same set of transcripts. The second scenario is that the same TE type is expressed, but from different genomic loci, consequently producing different transcripts. To explore this, we first needed to understand the relationship between TE type and expression level at the individual transcript level. We used bulk short-read RNA-seq data for each lineage with the long-read based hEDT reference, as the dynamic range of expression is higher for short-read data than long-read data^31^. In all four cell states, the expression levels of TE-free coding transcripts were significantly higher (T-test p-value < 0.05) than those of TE-containing coding transcripts (**Extended Data Fig. 9a)**. TEs inserted into the 5’ UTRs resulted in transcripts with the lowest expression levels while 3’ UTR insertion led to comparatively higher expression levels, but the levels were not as high as those of the TE-free transcripts.

We next investigated the aggregate expression patterns and found that the overall expression patterns for SINE, LTR, LINE, retroposon, and DNA coding transcripts were not substantially different across the four states (**Fig. 5a**). However, differential transcript expression between hPSC and the three differentiated cell states revealed that hPSC-to-endoderm differentiation induced the least number of transcript changes while mesoderm differentiation drove the largest coding transcript expression changes (**Extended Data Fig. 9b)**. Differentiation in all three lineages led to both upregulation and downregulation of TE-containing coding transcripts, suggesting that transcript regulation is locus-specific and that TE expression is not restricted to hPSCs. We checked the overlaps of the differentiation-induced differential transcripts and found that many transcripts were regulated in a lineage-specific manner (**Extended Data Fig. 9c)**. The aggregated expression of specific TE coding transcripts identified seven LTR TEs (LTR7Y, LTR7C, LTR7B, LTR7, HERVH, HERVH48 and HERVFH21) that were significantly downregulated in both ectoderm and mesoderm cells, and two LINE TEs (L1P4b_5end and L1HS_5end) that were substantially upregulated in ectoderm cells (**Fig. 5a**, **Extended Data Figs. 9d, e**). This indicates that although the TE types are similar in all three cell types, they originate from different transcripts.

**Fig. 5:**
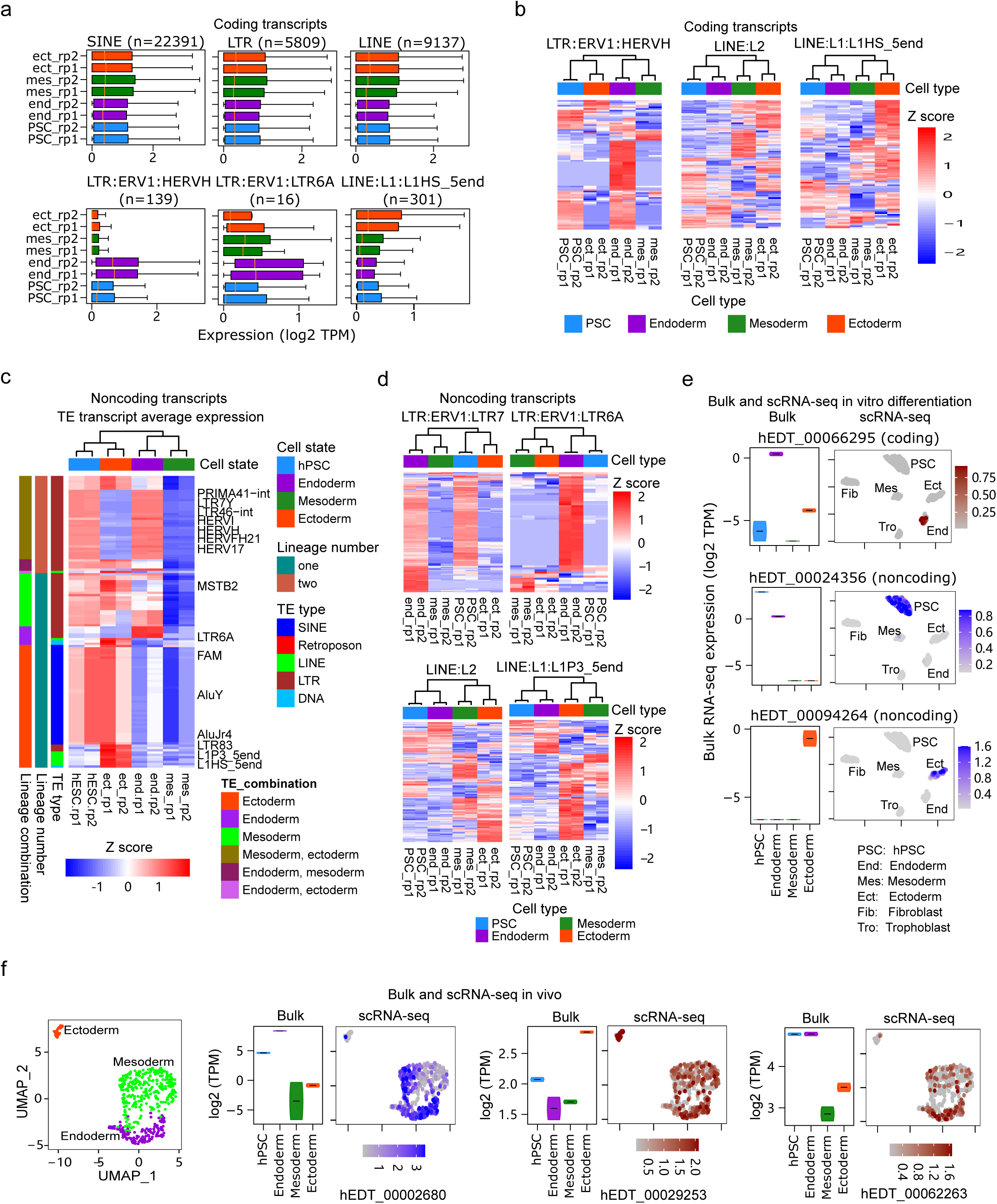
Dynamic expression patterns of TE-containing coding and noncoding transcripts across different cell states. **a**, Boxplots of the aggregate expression levels of coding transcripts containing specific TEs and TE types. **b**, Heatmap showing the expression changes of differentiation-induced differentially expressed coding transcripts containing the stated TEs. Each row of the heatmap represents a transcript containing the indicated type of TE (top label). **c**, Heatmap of aggregate changes in expression of noncoding transcripts containing different TEs. Each row represents the mean of all transcripts containing a TE-type that is significantly different (adjusted p-value < 0.05 and fold change >=2) between hPSC and any of the differentiated cell states. For each TE, the lineage(s) with significant difference, and the number of lineage(s) with significant differences are shown. **d**, Heatmap showing the change in expression levels of differentiation-induced differentially expressed noncoding transcripts containing the indicated TEs. Each row represents a TE-containing transcript that is differentially expressed between hPSC and any of the three differentiated states. Bulk RNA-seq (left) and scRNA-seq (right) expression patterns of selected TE-containing transcripts in *in vitro* (**e**) and *in vivo* (**f**) differentiation data.

We then examined the expression levels of the individual TE-containing coding transcripts. Surprisingly, the majority of the HERVH-containing differentially expressed coding transcripts were more highly expressed in endoderm cells but not in hPSCs (**Fig. 5b**). Different coding transcripts containing L2 and L1HS_5end were activated in different cell states, with activation of many L1HS_5end-containing coding transcripts in ectoderm cells. These data support the second of our scenarios, at least for coding transcripts, that although similar TE types are expressed, they are derived from different loci and are expressed in cell type-specific transcripts.

As noncoding transcripts are more likely to be expressed in a cell type-specific manner^31,59,60^, we expected that noncoding TE-containing transcripts would also be expressed from different transcript loci. However, in comparison to the coding transcripts (**Extended Data Fig. 9b**), fewer differentially expressed noncoding transcripts were induced by the differentiation process (**Extended Data Fig. 9g**), reflecting the lower number of noncoding transcripts in each cell type (**Fig. 2f**). Among the TE types, LTRs changed the most, with more LTR-containing noncoding transcripts downregulated in mesoderm cells. Also, many of the differentially expressed transcripts were regulated in a cell type-specific manner, and only a few differentially expressed transcripts were shared among cell states (**Extended Data Fig. 9h**). As for coding transcripts, in all cell states, TE-containing transcripts had lower overall expressions levels than TE-free transcripts (T-test p-value < 0.05) (**Extended Data Fig. 9i**). The comparison of the aggregate expression for noncoding transcripts containing specific TEs showed that 126 TEs were differentially expressed between the hPSC and at least one of the differentiated cell states (**Fig. 5c**). These TEs included multiple LTRs that are activated in hPSCs and endoderm cells.

Interestingly, several SINEs were also differentially expressed. LTR6A and a few other LTR TEs were also found to be enriched for endoderm cells. The analyses of the differentially expressed LTR7-containing transcripts showed that many of the transcripts are downregulated in ectoderm and mesoderm cells, whilst LTR6A-containing noncoding transcripts were up in endoderm (**Fig. 5d**). Also, specific LINE-containing transcripts were specifically expressed in different lineages. Interestingly, half of the TEs with significant expression differences also had significantly different splicing frequencies (**Extended Data Fig. 9j**). These data support the model that similar TE types are expressed from different transcripts in different cell types.

We next investigated the expression dynamics of TE-containing coding transcripts at the single-cell level using scRNA-seq data from *in vitro* differentiated germ lineage cells^51^. The scRNA-seq data recaptured the higher expression of multiple TEs in ectoderm cells and downregulation of some LTRs such as LTR6A, LTR7, LTR7C, and HERVH in ectoderm and mesoderm cells (**Extended Data** Fig. 10a). The scRNA-seq data also revealed that the expression differences were not obvious at the level of TE types but became distinct at individual transcripts (**Fig. 5e, Extended Data Fig. 10b)**. Also, single-cell analyses of the noncoding transcripts were consistent with the observations from bulk data, and revealed upregulation of multiple TEs in ectoderm cells (**Extended Data Figs. 10a, b)**. Several LTR TEs were downregulated in ectoderm and mesoderm, while LTR6A was enriched in endoderm cells (**Fig. 5e, Extended Data Figs. 10c-d**). As with bulk data, lineage-specific expression became more obvious when TEs were considered at the transcript level (**Fig. 5e, Extended Data Fig. 10d**). These results demonstrated that the observations in bulk RNA-seq were recaptured in single cells.

To further elaborate on the expression dynamics of TE-containing transcripts we extended our analysis to scRNA-seq from *in vivo* gastrulation data^61^ (**Fig. 5f, Extended Data Fig. 11a**). As expected, ectoderm cells were marked by more TE-containing coding and noncoding transcripts (**Extended Data Figs. 11a, b)**. Indeed, compared to other cell types, more ectoderm-expressed transcripts were found in almost all of the TE types in both coding and noncoding transcripts (**Extended Data Figs. 11c)**. Also, many transcripts that were identified in the *in vitro* data were recaptured in the *in vivo* data (**Fig. 5d, Extended Data Fig. 11d**), indicating that *in vitro* observations reflected *in vivo* phenomena. Taken together, these data revealed that whilst similar TE types are expressed in all cell types, they originate from multiple lineage-specific transcripts, suggesting that TEs of the same type from multiple loci are independently regulated.

To investigate the regulation of hEDTs, we checked the enrichment of transcription factor binding motifs in the core promoter (200 bp upstream of the TSS) for the assembled transcripts that were differentially expressed across different differentiation courses. We used expressed transcripts without significant differentiation-induced expression changes as the control. As expected for endoderm, motifs for pluripotent transcription factors (TFs) like POU5F1 (OCT4) were enriched in transcripts with higher expression in PSC (down in endoderm), while the motifs for endoderm marker EOMES and GATA4 were in transcripts upregulated in endoderm (**Extended Data Fig. 12**). Some TFs like MAfF were found to be consistently enriched in transcripts that were downregulated during differentiation. This data suggest that the lineages had both shared and unique TFs activated during differentiation. Next, we checked the enrichments of RNA-binding protein motifs in the differentially expressed transcripts using transcripts without differentiation-induced expression changes as the background. However, a number of RNA-binding proteins (RBPs) were enriched such as HNRNPK, PCBP2 and PCBP1 were enriched in both upregulated and downregulated transcripts (**Extended Data Fig. 13**), suggesting that the RBPs might be involved in cell fate determination without any particular fate preference. These analyses highlighted complex transcript expression regulation during differentiation.

### TE-type switching of transcripts during differentiation into somatic cells

As the same TE types are expressed from different transcripts, we next took advantage of differentiation time course data to investigate how the expressions of TE-containing transcripts vary during the endoderm differentiation process. As many LTR6A and HERVH-containing transcripts are specifically expressed in the endoderm or hPSCs (**Figs. 5b, d)**, we focused our initial analyses on the transcripts containing these two TEs. The analyses of the time-course bulk RNA-seq data showed that the activation of LTR6A noncoding transcripts happened at 72 hours in the differentiation process (**Figs. 6a, b)**. HERVH-containing noncoding transcripts with higher endoderm expression were upregulated as early as 24 hours but did not reach maximum expression until 72 hours. Conversely, HERVH-containing noncoding transcripts with high expression in hPSCs were downregulated in endoderm at as early as 12 hours and were essentially undetectable at 72 hours of differentiation to endoderm. The situation was similar for HERVH-containing coding transcripts (**Figs. 6a, Extended Data Fig. 14a**). These results demonstrate how HERVH and LTR6A can both be expressed in hPSCs and endoderm, but these TE sequences are contained in lineage-specific transcripts.

**Fig. 6:**
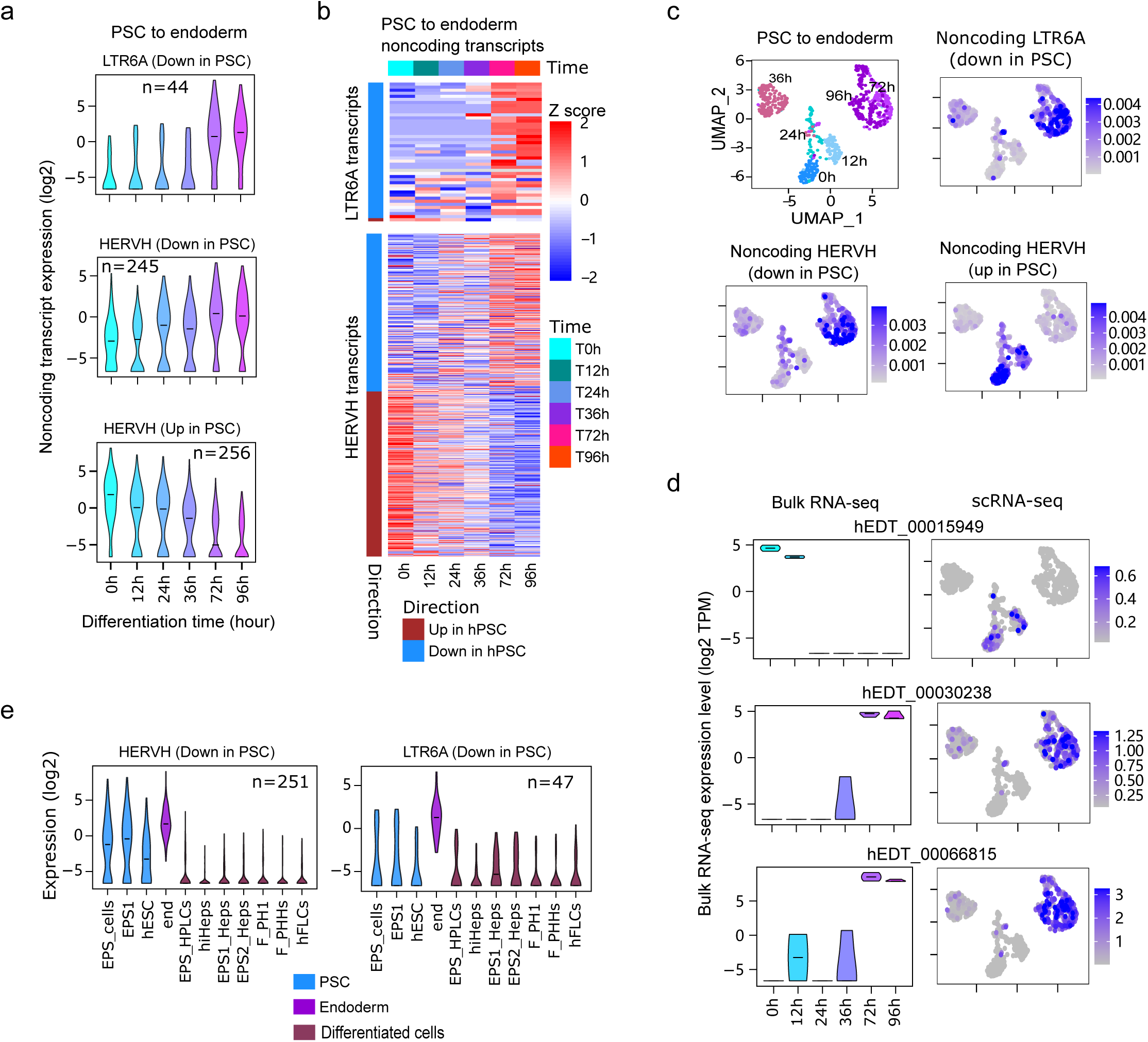
Expression changes of TE-containing transcripts in endoderm lineage. **a**, Violin plots of bulk RNA-seq expression levels of HERVH and LTR6A-containing noncoding transcripts during differentiation to endoderm. The data were from GSE75748. **b**, Heatmap showing the expression dynamics of HERVH and LTR6A noncoding transcripts differentially expressed in endoderm lineage. **c**, UMAP showing the single-cell expression data for a hPSC to endoderm time course. Hours of differentiation are shown. The three other UMAPs show example aggregate scores for transcripts containing the indicated TE types that are down-regulated in hPSCs or up-regulated in hPSCs. **d**, Expression dynamics in bulk RNA-seq (left) and scRNA-seq (right) of selected transcripts. **e**, Expression dynamics of endoderm-upregulated HERVH and LTR6A-containing noncoding transcripts in hepatocyte-related cells, using transcripts containing the indicated TE types, that are defined as down-regulated in hPSCs. (EPS: Extended pluripotent stem cells; EPS1: Stage 1 EPS; EPS2: Stage 2 EPS; hESC: Human ESC (H1); HPLC: Hepatic progenitor -like cells; EPS_HPLCs: EPS-derived HPLCs; hiHEPs: iPSC-derived hepatocytes; EPS1_Heps: EPS1-derived hepatocytes; EPS2_Heps: EPS2-derived hepatocytes; F_PH1: Fetal primary hepatocytes; F_PHHs: Fresh primary human hepatocytes; hFLC: Human fetal liver cells).

Next, we analyzed scRNA-seq data of time-course hPSC-to-endoderm differentiation. Cell clustering by identity showed that the cells formed three distinct clusters representing early (0h to 24h) middle (36h) and late (72h and 96h) stages of differentiation (**Fig. 6c**). In agreement with the bulk RNA-seq analysis, endoderm-upregulated LTR6A, and HERVH-containing noncoding transcripts had high expression only at 72- and 96-hour of differentiation. In contrast, endoderm-downregulated noncoding transcripts were shut down at 12 hours of differentiation (**Fig. 6d**). These data showed that the timing for the activation and downregulation of TE transcripts varied during differentiation.

To further investigate if the expression of the endoderm-induced upregulated transcripts were maintained in terminally differentiated somatic cells, we first checked their expression in an array of hepatocyte-related samples. Surprisingly, the expression of noncoding transcripts with endoderm-induced upregulation was not sustained in hepatocyte-related terminally differentiated somatic cells (**Fig. 6e, Extended Data Fig. 14b, c**). Similar observations were found for HERVH-containing endoderm-upregulated coding transcripts (**Extended Data Fig. 14d**). Importantly, the expression of endoderm-specific HERVH-containing noncoding transcripts was not maintained in endoderm-derived terminally differentiated somatic cells (**Extended Data Fig. 15**). Conversely to HERVH, the expression of noncoding transcripts containing L1P1_orf2 and LTR that were specifically expressed in the ectoderm were maintained in multiple ectoderm-derived somatic cells, such as neurons (**Extended Data Fig. 15**). Similarly, the expression of many ectoderm-activated coding transcripts containing L2, L1P1_orf2 and LTR remained active in ectoderm-derived terminally differentiated somatic tissues. In summary, these results demonstrate how TE types can remain active across cell types, but they are often expressed from different transcripts in distinct tissues.

### TE presence influences transcript subcellular localization

Previous studies have shown that TE-containing transcripts tend to localize in the nucleus^62,63^. To this end, we generated RNA-seq data for subcellular fractions of hPSCs, mesoderm, endoderm, and ectoderm differentiated cells for the nucleus and cytoplasm. Western blots for proteins known to localize to the nucleus and cytoplasm confirmed that the intended cellular fractions were purified (**Extended Data Fig. 16a**). We confirmed our previous observation that both coding and noncoding TE-containing transcripts preferentially localize to the nucleus in hPSCs^27^ (**Fig. 7a**). This pattern was the same for mesoderm and endoderm. Surprisingly, this was not the case for ectoderm, where the opposite pattern was seen: TE-containing transcripts were instead more likely to localize in the cytoplasm. Indeed, analyses of TE types revealed that they all tended to preferentially localize to the cytoplasm in ectoderm cells, and to the nucleus in all other cell states (**Extended Data Fig. 16b**). Greater details were revealed at the transcript level. For example, LTR-containing noncoding transcripts tended to have different cytoplasm/nucleus enrichment when compared to other TE types in PSC and endoderm (**Extended Data Fig. 16c**).

**Fig. 7:**
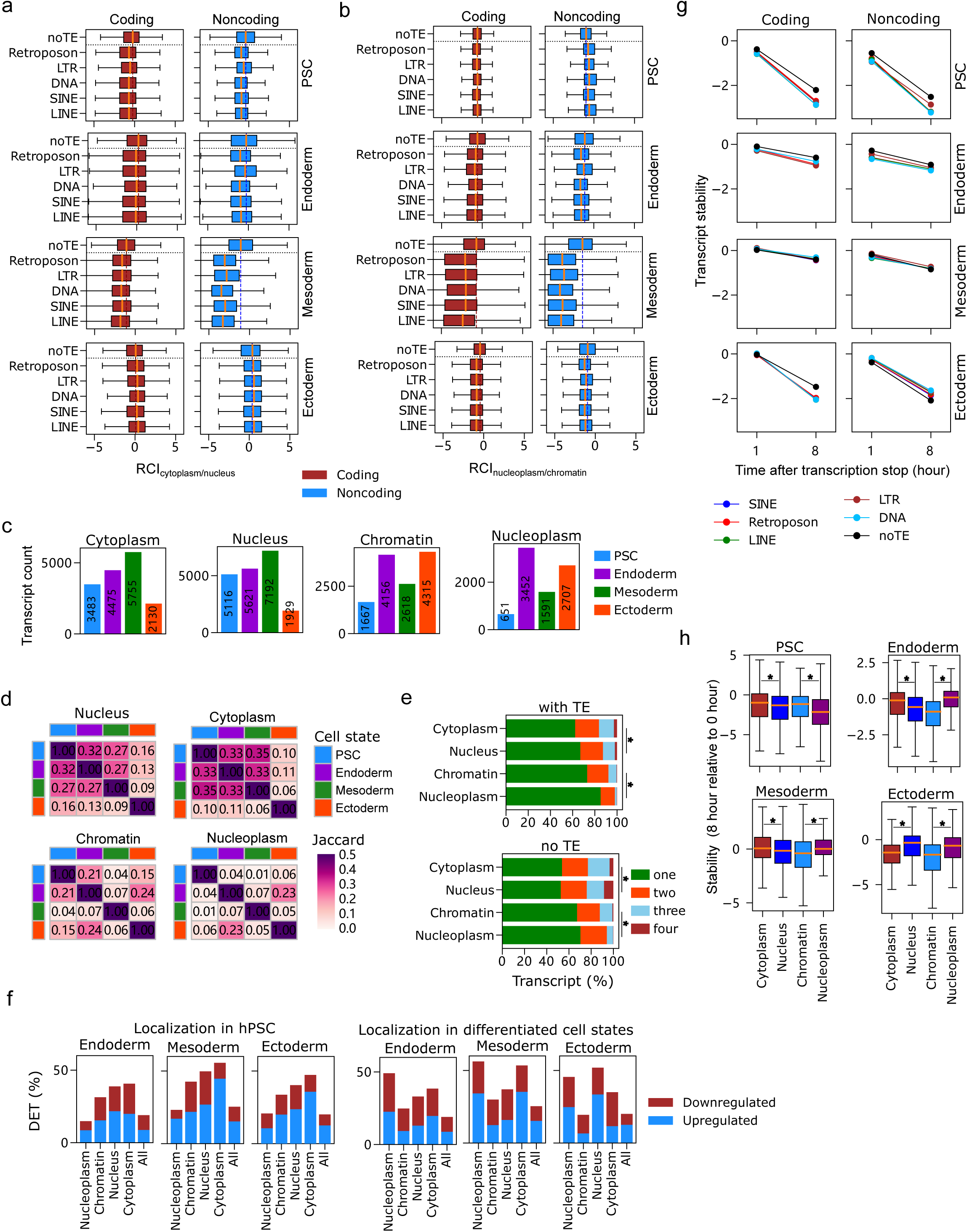
Subcellular localization and stability of TE-containing transcripts. **a**, Cytoplasm/nucleus relative concentration index (RCI). RCI is the log2-transformed relative expression in two tissues. **b**, Nucleoplasm/chromatin RCI. **c**, Bar charts showing the number of transcripts localized to different subcellular structures in different cell states using individual cell state assemblies. **d**, Heatmap showing subcellular localization consistency of hEDT transcripts localized to different structure across cell states. **e**, Stacked bars showing the distribution of the number of cell states in which transcripts are localized to a subcellular structure. (* represents Chi-square significance at p value < 0.05). **f**, The proportion of localized transcripts that were differentially expressed during differentiation process. The left charts show the localization in hPSC while the right charts show the localization in differentiated cell states. The distributions for all localizations were significantly different from the distribution of all hEDT set (Chi-square p value < 0.05). **g**, Line plots showing the transcript stability of TE-containing transcripts, using time from actinomycin D treated cells. The expression levels at time 1 and 8 hours, relative to the expression at 0 hour are shown for each TE type across different cell states. **h**, Boxplots showing the stability of transcripts that are localized to different subcellular structures across cell states (* represents Mann-Whitney test significance at p value < 0.05).

Noncoding RNAs that localize to the nucleus are often recruited to chromatin^64,65^. Hence, we performed RNA-seq on the nucleoplasm and chromatin fractions to understand the sub-nuclear localization of TE-containing transcripts (**Extended Data Fig. 16a**). TE-containing hPSC transcripts were reduced in the chromatin compartment, versus the nucleoplasm (**Fig. 7b)**. On the contrary, the TE containing transcripts of endoderm, mesoderm and ectoderm tended to be more enriched in the chromatin fraction (**Fig. 7b, Extended Data Fig. 16d)**. Transcript-level analyses of nucleoplasm/chromatin enrichment revealed heterogeneity across coding potentials, cell states and TE types (**Extended Data Fig. 16e)**. These data reveal that the sub-cellular and sub-nuclear localization of TE-containing transcripts is cell type-specific, and retention of TE-containing transcripts in the nucleus is not a general feature.

To further investigate the subcellular transcript localization, we compared the expression levels of different transcripts to obtain transcripts enriched in specific sub-cellular fractions (see Supplementary methods). Pairwise comparisons of nucleus-cytoplasm and chromatin-nucleoplasm expression patterns identified transcripts that were enriched in specific sub-cellular fractions (**Extended Data Fig. 17)**. The number of transcripts localized to the sub-cellular compartments varied across cell types (**Fig. 7c**). Importantly, compared to the differentiated cells, hPSCs had fewer transcripts differentially localized to the nucleoplasm and chromatin. The distribution of coding and noncoding transcripts revealed both cell state and location-specific differences. Specifically, while nucleoplasm-localized transcripts had the highest noncoding proportion in hPSC, chromatin-localized transcripts had the highest noncoding proportion in differentiated cells (**Extended Data Fig. 18a**). Similarly, the distribution of TE-containing transcripts varied across cell types, subcellular location, and protein-coding ability (**Extended Data Fig. 18b**). This effect was also transcript-specific, as few transcripts overlapped in the distinct sub-cellular compartments between the different cell types (**Extended Data Fig. 19**). Indeed, using a Jaccard similarity measure, transcripts that localized to the nucleus or cytoplasm were more consistent in hPSC, endoderm and mesoderm but more divergent in ectoderm (**Figs. 7d**). Interestingly, we found that the distribution of transcripts based on the number of cells in which they were localized varied across subcellular structures (**Fig. 7e**). The most unique chromatin localization was found in the mesoderm (**Figs. 7d**). Overall, nucleoplasm localization was the least consistent across the cell states. To further explore the relationship between transcript subcellular localization and cell state conversion, we checked the proportion of differentiation-induced DETs among the transcripts localized to various subcellular structures across cell states and found that subcellular localization significantly influenced the proportion of DETs (**Fig. 7f**), as the distribution of DETs were significantly different in localized transcripts compared to overall transcripts (Chi-square p-value < 0.05). For hPSCs, nucleoplasm-enriched transcripts had the lowest proportion of DETs. For differentiated cells, however, the lowest proportion of DETs was found in chromatin-localized transcripts. These data indicate that the localization of TE-containing transcripts is cell type-specific. Overall, compared to the TE-free transcripts, we found that TE-containing transcripts were more cell type-specific (**Fig. 7e**), with hPSCs having fewer chromatin-enriched TE transcripts.

An important question was how the localization of the transcript to different subcellular structure was established. We hypothesized that the activities of RBPs might contribute to the transcript localization. Consequently, we checked the RBP enrichment of localized transcripts with expressed transcripts with no significant bias for each cell state. Interestingly, we found multiple RBPs that were enriched in transcripts localized to specific subcellular structures (**Extended Data Fig. 20**). Interestingly, RBPs that were associated with chromatin localization such as SAMD4A, PCBP2 and LIN28A were also found to be associated with nucleus localization across different cell states. Also, some RBPs like PABPN1 were found to be associated with cytoplasm and nucleoplasm localization across cell states. These results suggest that chromatin enrichment might contribute to nucleus enrichment, and that RBPs might be implicated in transcript localization to different subcellular structures.

### TE presence influences transcript stability

As subcellular localization can influence degradation, we wondered if this would impact transcript half-life in a TE, sub-cellular localization, and cell state-dependent manner. We performed RNA-seq in differentiated cells treated with Actinomycin D to block transcription for 1 and 8 hours and measured the transcript levels relative to the 0-hour time-point. Surprisingly, there were divergent patterns of transcript half-life across TE types, coding potentials, and cell states (**Fig. 7g**). The lowest stability was found in hPSCs and endoderm. Interestingly, the impact of TE presence was also dynamic. While the TE-free transcripts were more stable in hPSC, ectoderm, and endoderm, this was not the case in mesoderm (**Fig. 7g, Extended Data Fig. 21a, b**). TE-containing ectoderm transcripts also contrasted with hPSCs and endoderm, as whilst TE-containing coding transcripts were less stable, TE-containing noncoding transcripts were more stable than TE-free transcripts (**Fig. 7g**, **Extended Data Fig. 21a, b**). These revealed that transcript stability varied across cell states and transcript coding ability.

Finally, we checked if TE presence had any substantial influence on the stability of transcripts with differential subcellular localization. The stability at 8 hours after transcription stop showed differences across transcripts localized to different subcellular structures (**Fig. 7h**). While nuclear-localized transcripts were more stable than cytoplasm-localized transcripts in the ectoderm, the opposite was true in the other cell states. Conversely, chromatin-localized transcripts were more stable than nucleoplasm-localized transcripts in hPSCs, but the nucleoplasm-localized transcripts were more stable than chromatin-localized transcripts in differentiated cells. Interestingly, some differences in stability can be observed as early as 1 hour after transcription stop (**Extended Data Fig. 21c**). Moreover, the comparison of stabilities of TE-free transcripts to those containing TEs identified multiple TEs associated with significant differences in stability after 8 hours (**Extended Data Fig. 21d**). In hPSCs, TE presence mostly led to lower stability. In differentiated samples, mosaic patterns were found. For example, TE-containing transcripts localized to cytoplasm and nucleoplasm in mesoderm were more stable, along with cytoplasm-localized transcripts in ectoderm. These data suggest that TE presence might have different effects on transcript localization and stability across different cell states. Overall, the data suggest that stability and subcellular localization of TE-containing transcripts are dynamic across different cell state conversions

## Discussion

TEs are difficult to place in their genomic context due to their repetitive nature. However, understanding expression dynamics would substantially benefit from robust TE-containing transcript assemblies^27,36^. Here, we used long-read RNA-seq technology to generate an accurate TE-containing transcript assembly^30,31^. These assembled transcripts enabled the accurate placement of TE sequences into transcripts in hPSCs and *in vitro-*derived germ layers. Our results showed that TEs are often expressed alongside unique genomic sequences. This partial overlap suggests that the TEs might have been inserted into a transcribed unit, relying on other transcriptional machinery^1^. This possibility is supported by the evolutionary suppression of TE-sequences at the TSS and the relative overrepresentation inside the 3’UTRs of coding transcripts.

This study showed widespread and dynamic TE expression in human early development, although the frequencies and expression levels of different TE types vary between cell types. Multiple studies have shown the overexpression of TEs in hPSCs^10,11,66,67^. However, here we find that the expression of TE sequences is not restricted to hPSCs, and multiple TEs are expressed in both hPSCs and differentiated germ layers, Interestingly, although the same TE types are expressed, they are found in different transcripts. The ectoderm was particularly rich for TE-containing transcripts, and the expression of many TEs in ectoderm is sustained in the later stage of development, consistent with the reports of TE activity in terminally differentiated ectoderm-derived cells and tissues^34,68–70^. The exemptions to higher TE activities in ectoderm involved HERVH, LTR7, LTR6A, and several other LTRs that have previously been reported to be enriched in hPSCs^11,57^. In addition to expression in hPSCs, HERVH is also expressed in endoderm, and some HERVH-containing transcripts were higher in endoderm while others were higher in hPSCs.

Curiously, many of the endoderm-expressed HERVH transcripts were not expressed in endoderm-derived somatic cells, again revealing transcript switching.Consistent with previous studies ^27,71,72^, we found that TE presence leads to lower expression levels, probably due to the activities of RNA-binding proteins^16,72^. Indeed, SINE elements are overrepresented in 3’ UTRs, and STAU1 promotes mRNA decay by targeting those Alu elements^16^. This indicates that the meta-analysis of TEs is limited, and TE sequences should be considered in their transcript context. From a biological perspective, it also suggests that cell-state specific regulation of TE expression occurs at the transcriptional stage or post-transcriptional stage. Interestingly, we found that TEs tend to be enriched in the chromatin fraction of the differentiated cells, suggesting that TE enrichment on chromatin might be associated with chromatin changes which might induce or regulate cell fate transitions. Whether all the TE expression changes are drivers or footprints of cell differentiation require further detailed studies. However, a number of studies have demonstrated the roles of TE-containing transcripts in pluripotency^73–75^, cardiomyocyte development^76^ and neurogenesis^77–79^. Taken together, this study reveals that TE-containing transcripts are highly dynamic in human early development, and the dynamics contribute to the cell state transitions by regulating chromatin structure in differentiating cells.

## Acknowledgments

We acknowledge the assistance of SUSTech Core Research Facilities. Funding was from the National Natural Science Foundation of China (32270597, 31850410486) the Science Technology and Innovation Commission of Shenzhen (RCBS20221008093109033), and the Guangdong Basic and Applied Basic Research Foundation (2023A1515111170).

## Accession data

Sequencing data from this study was deposited in the Gene Expression Omnibus with accession numbers GSE269270, GSE269272, GSE269273, and GSE269274.

## Author contributions

I.A.B. planned and designed the study, performed most of the bioinformatic analyses, drafted the paper, and coordinated the experimental work. X.F. and G.M. performed the majority of the experiments. Y.L., M.T.A., and X.M. assisted with analysis. All authors assisted in writing the manuscript. A.P.H. designed the study, revised the manuscript, and supervised and funded the study.

## Methods

The detailed experimental procedure and bioinformatics pipelines for this study are presented in the Supplementary Methods.

