## Supplementary Figures for "Transposable Element Expression and Sub-cellular Dynamics During hPSC Differentiation to Endoderm, Mesoderm, and Ectoderm Lineages"

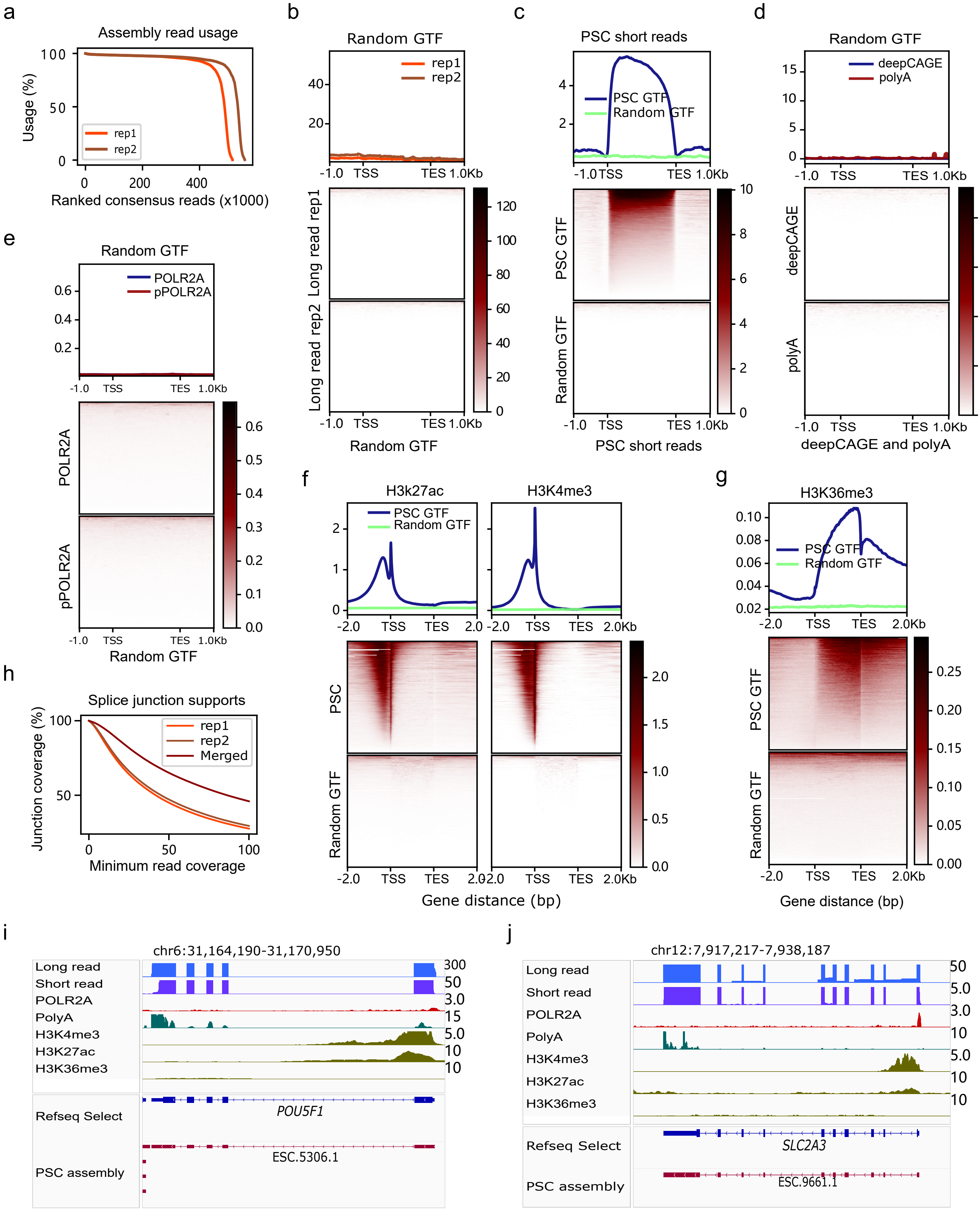


Extended Data Fig. 1

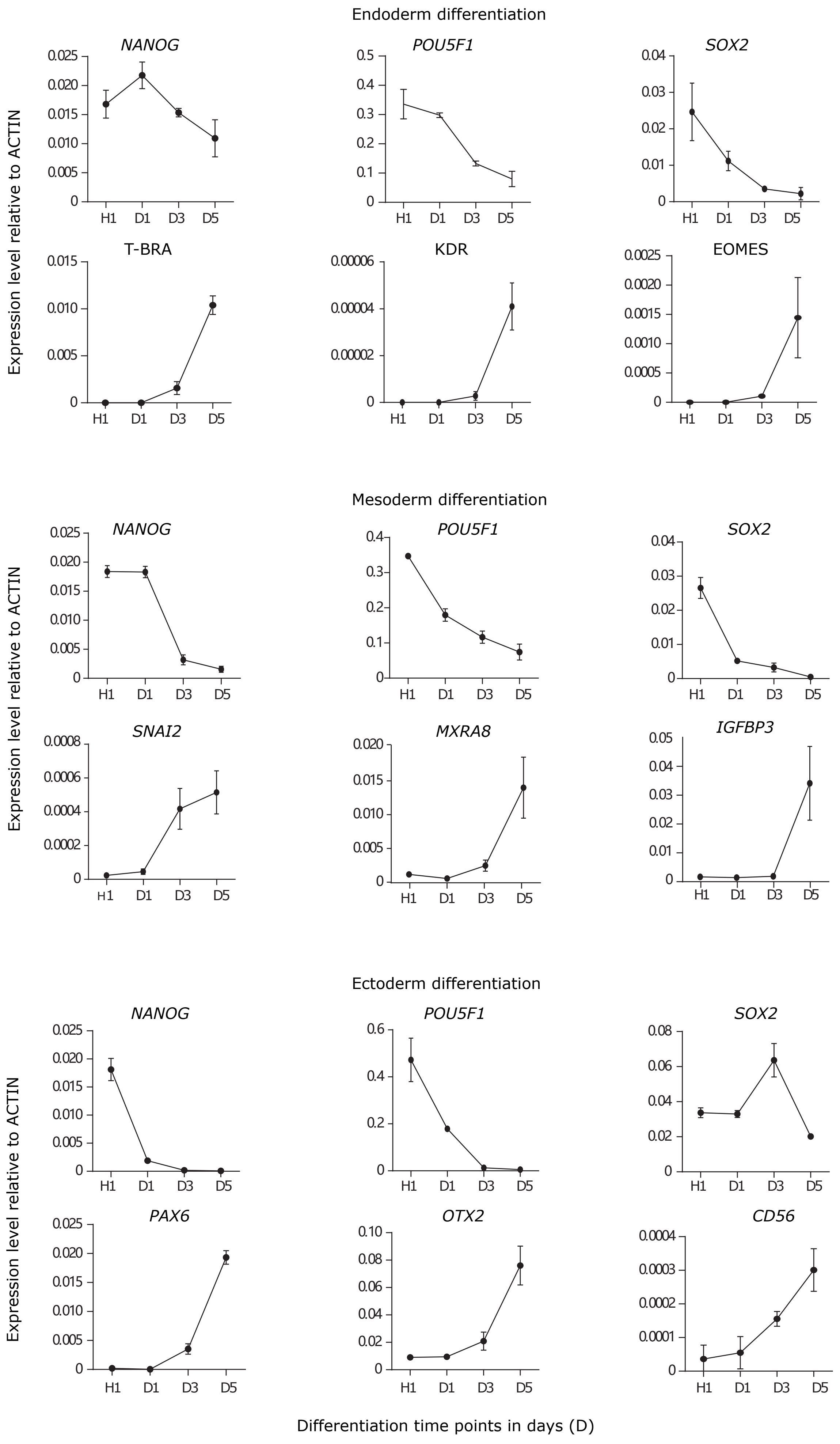


Extended Data Fig. 2

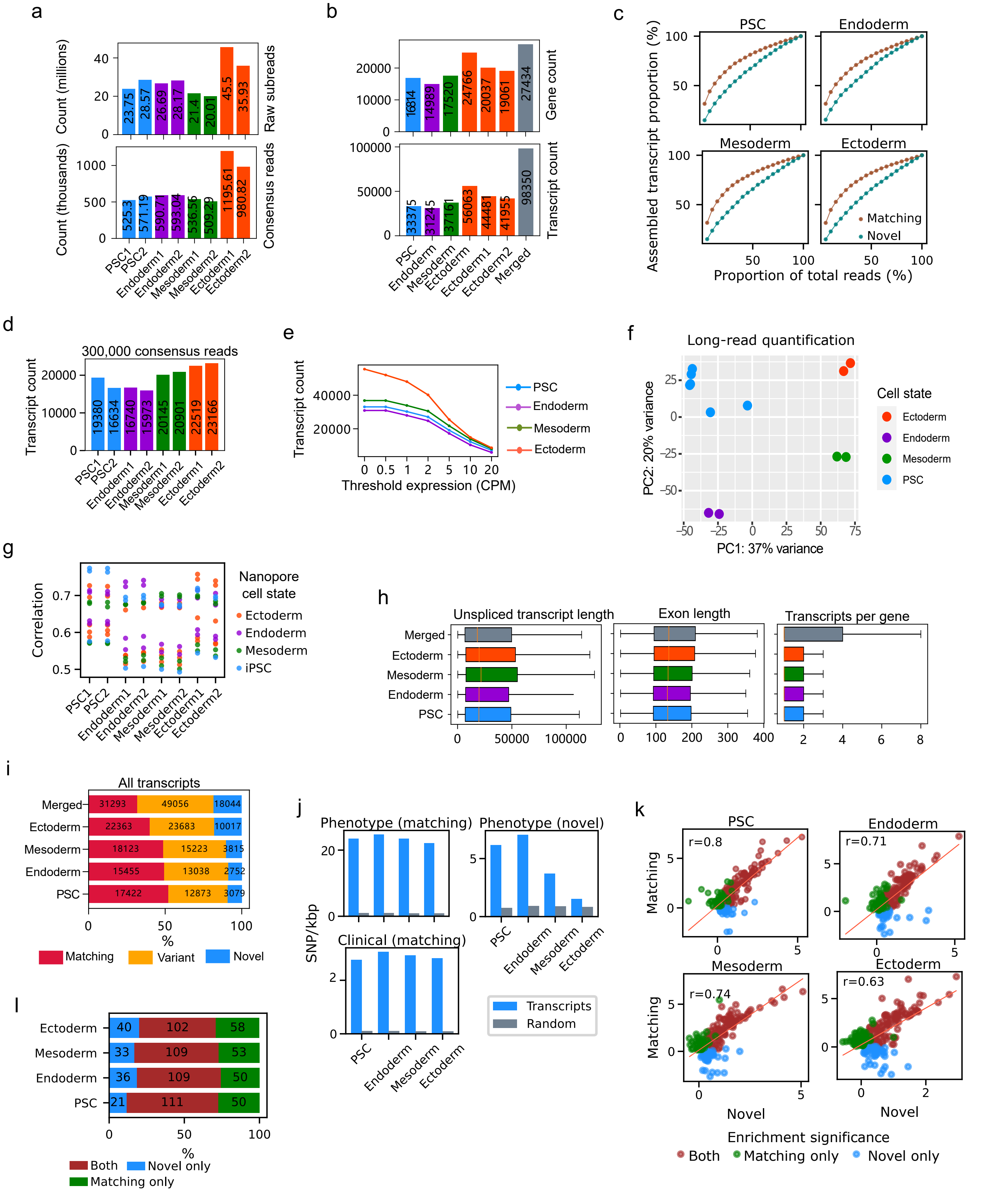


Extended Data Fig. 3

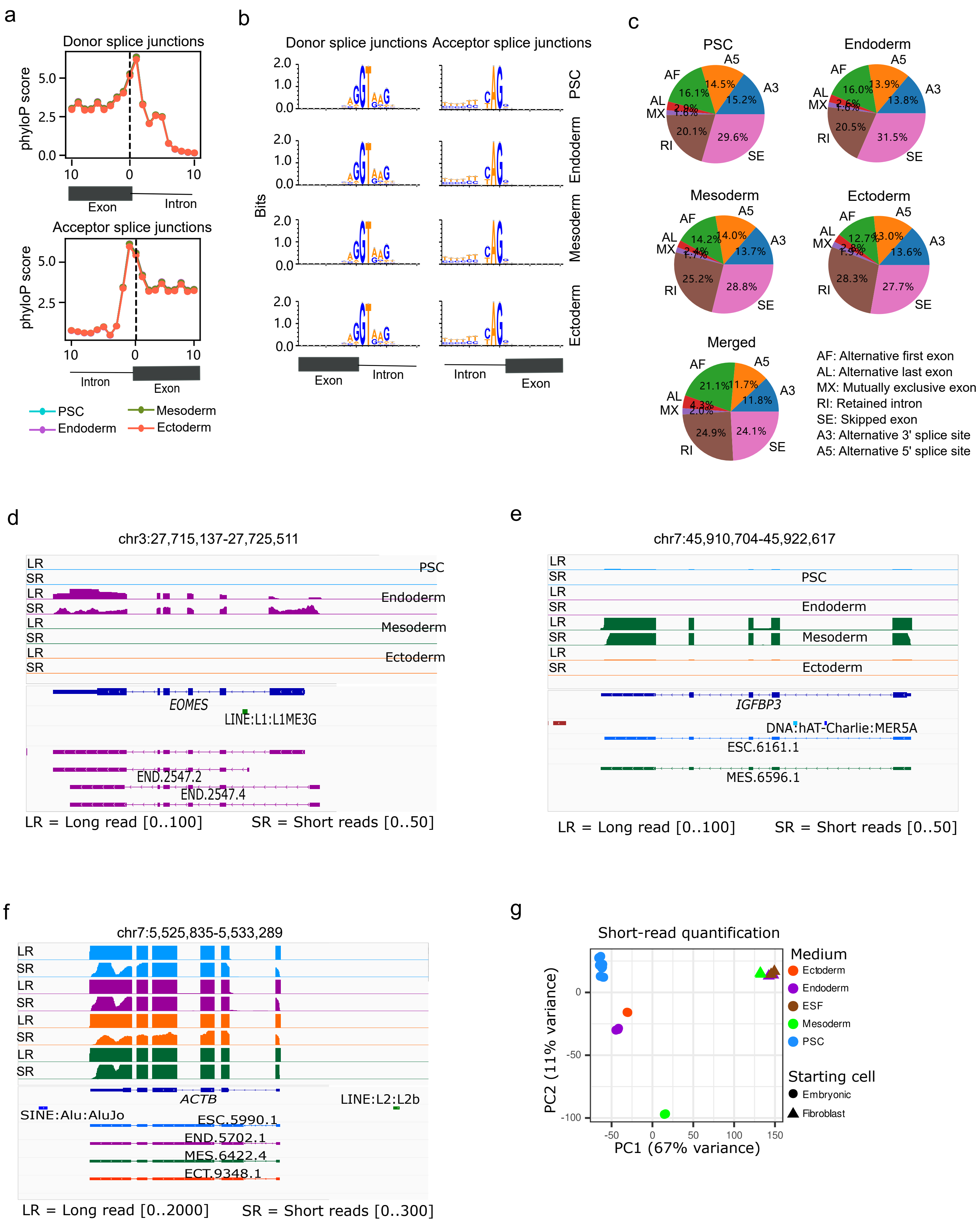


Extended Data Fig. 4

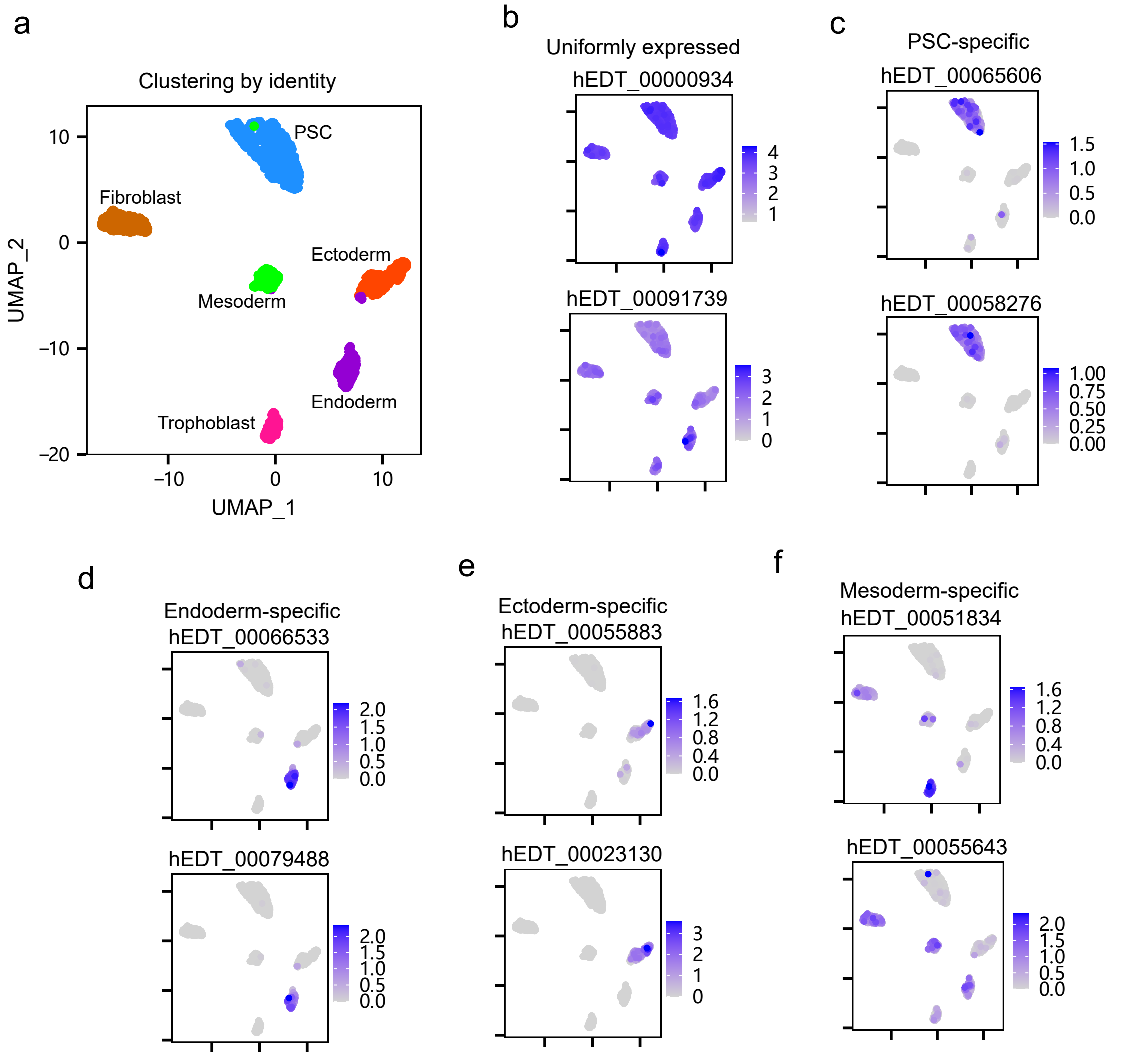


Extended Data Fig. 5

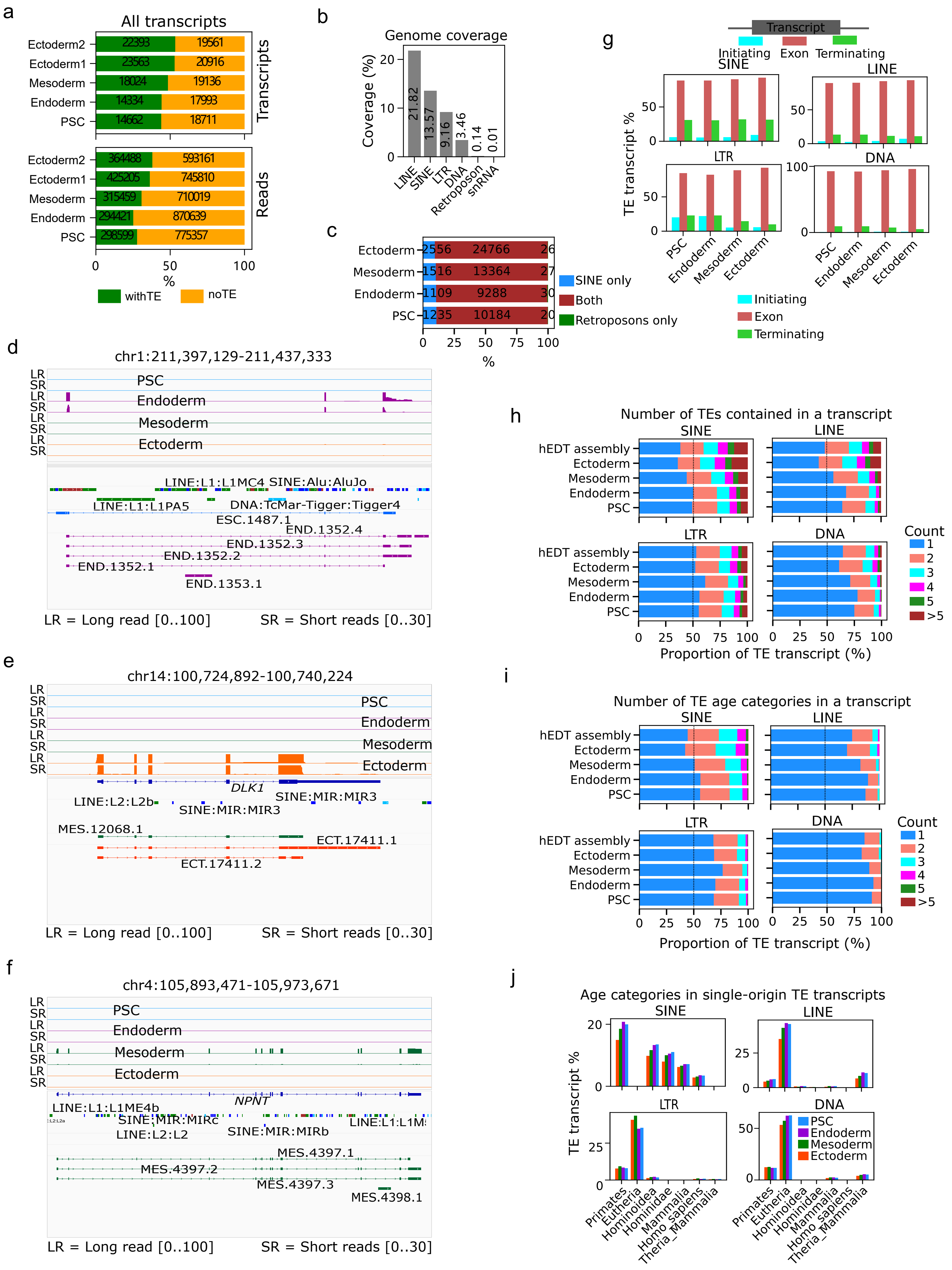


Extended Data Fig. 6

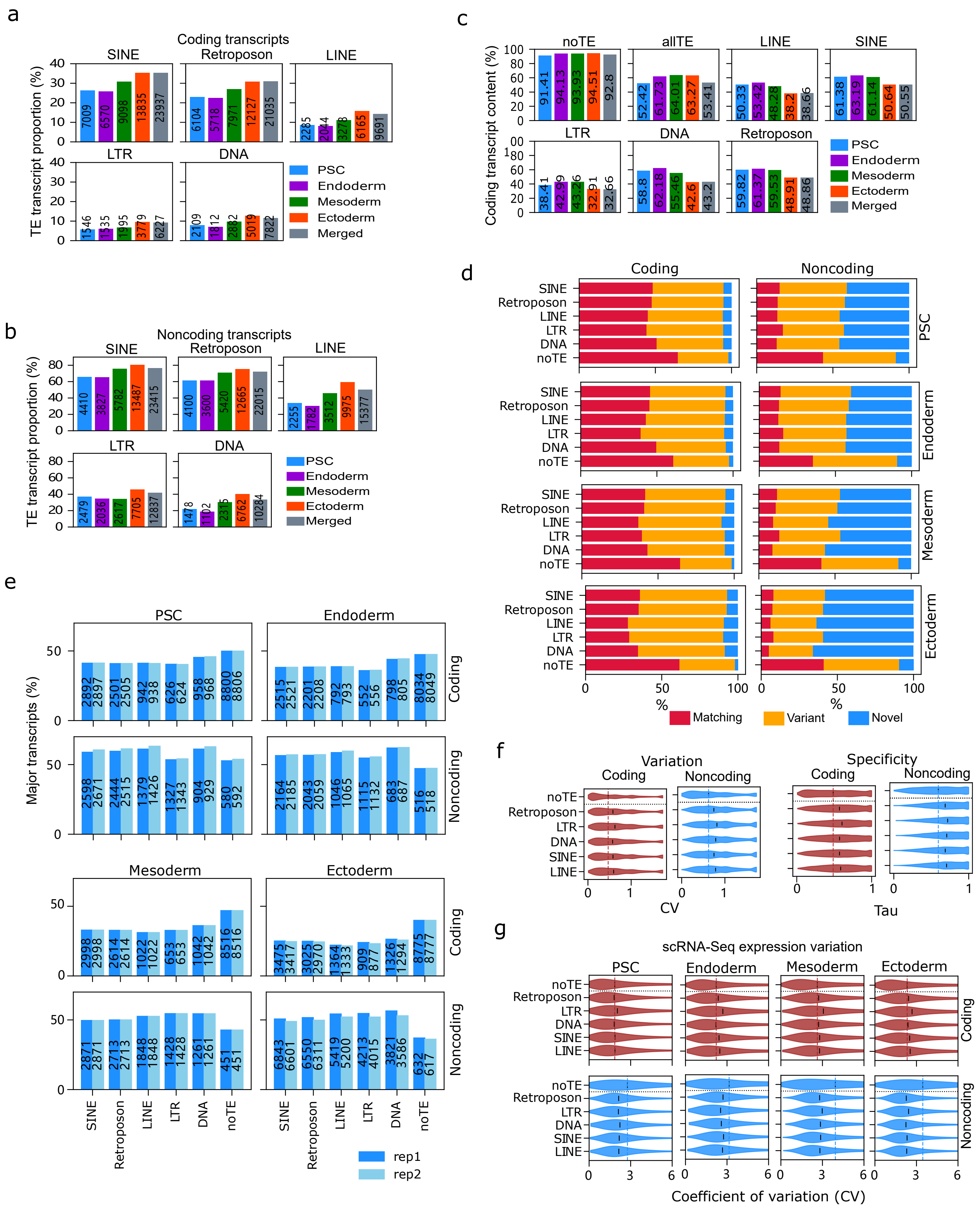

Extended Data Fig. 7

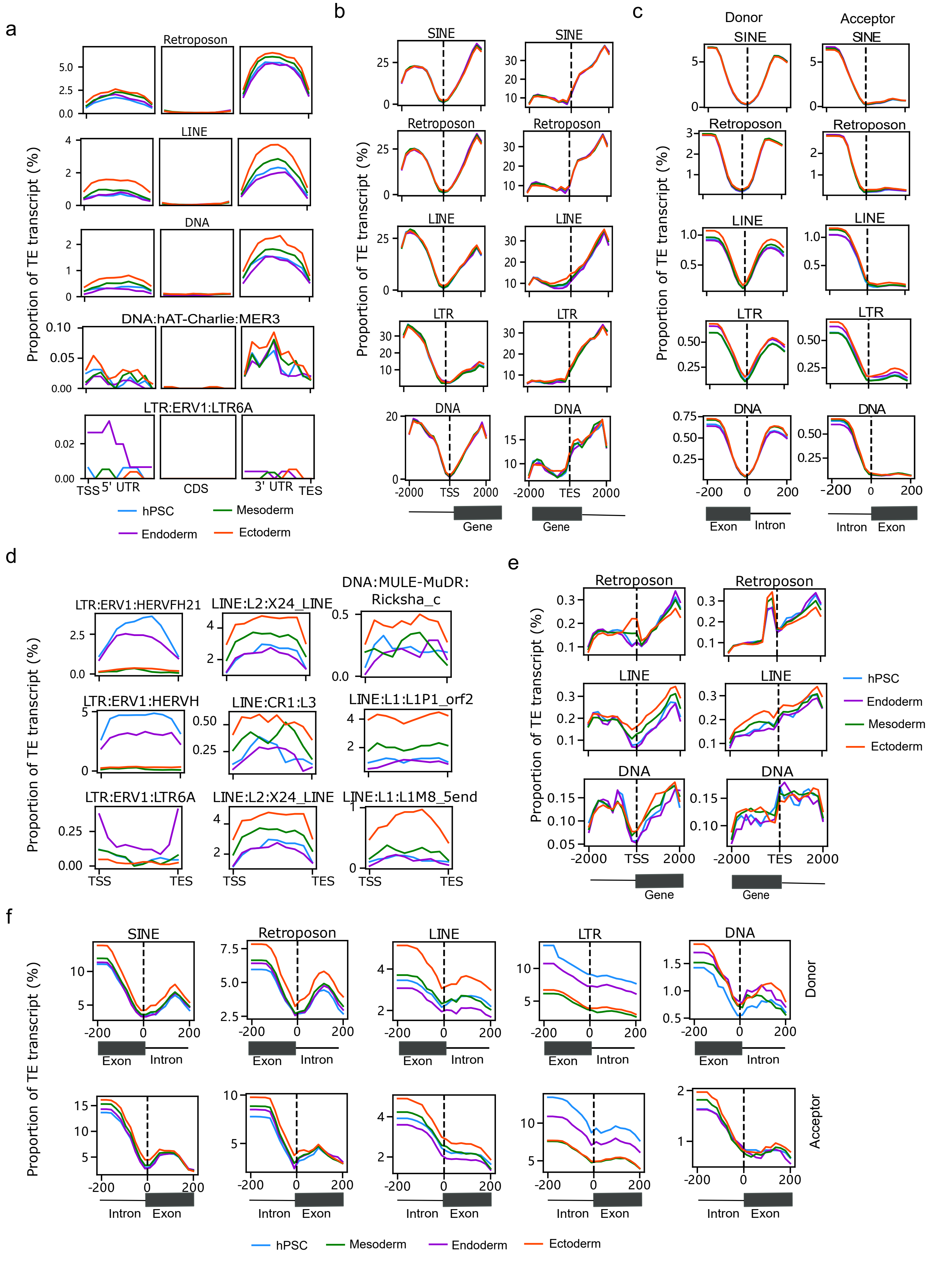


Extended Data Fig. 8

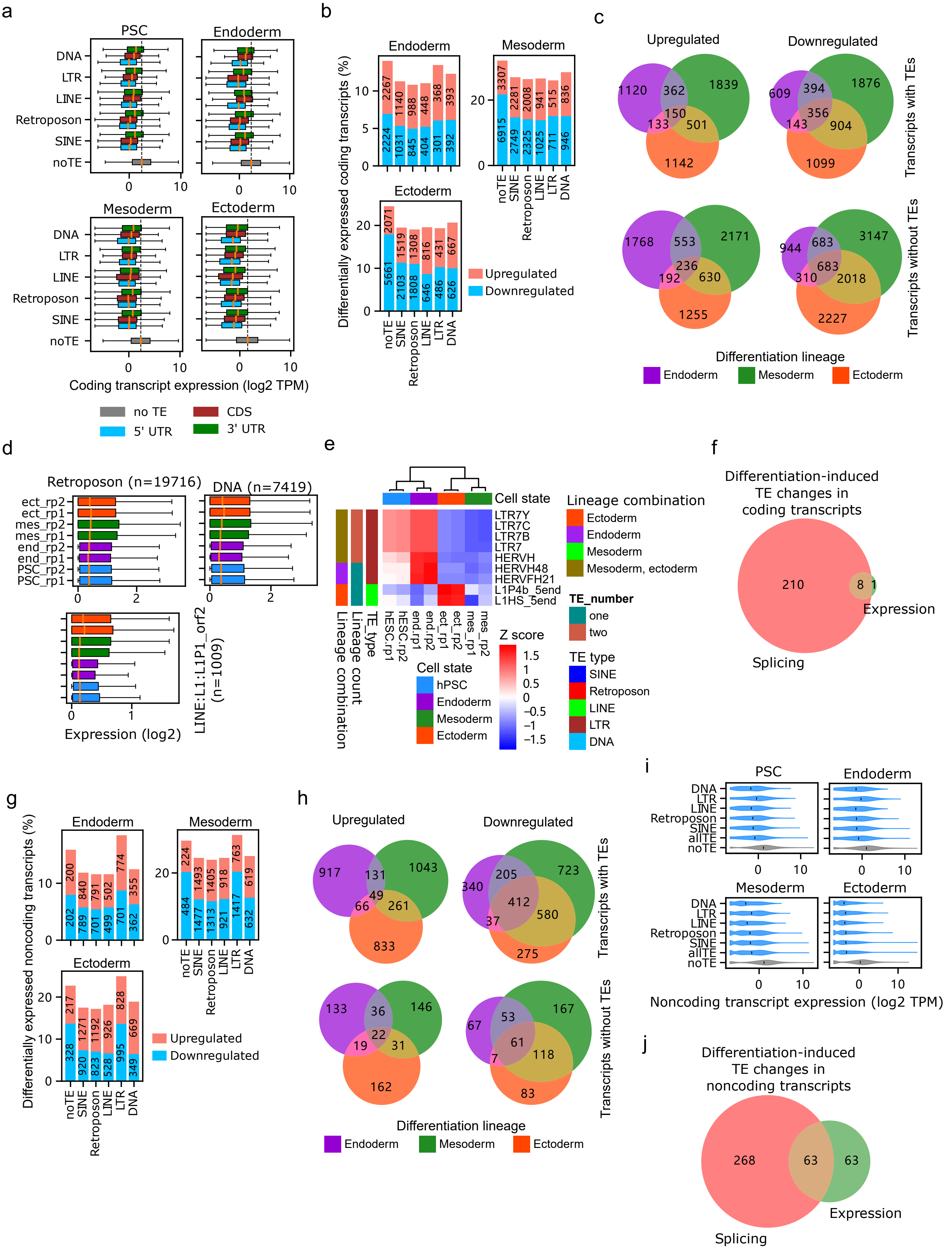


Extended Data Fig. 9

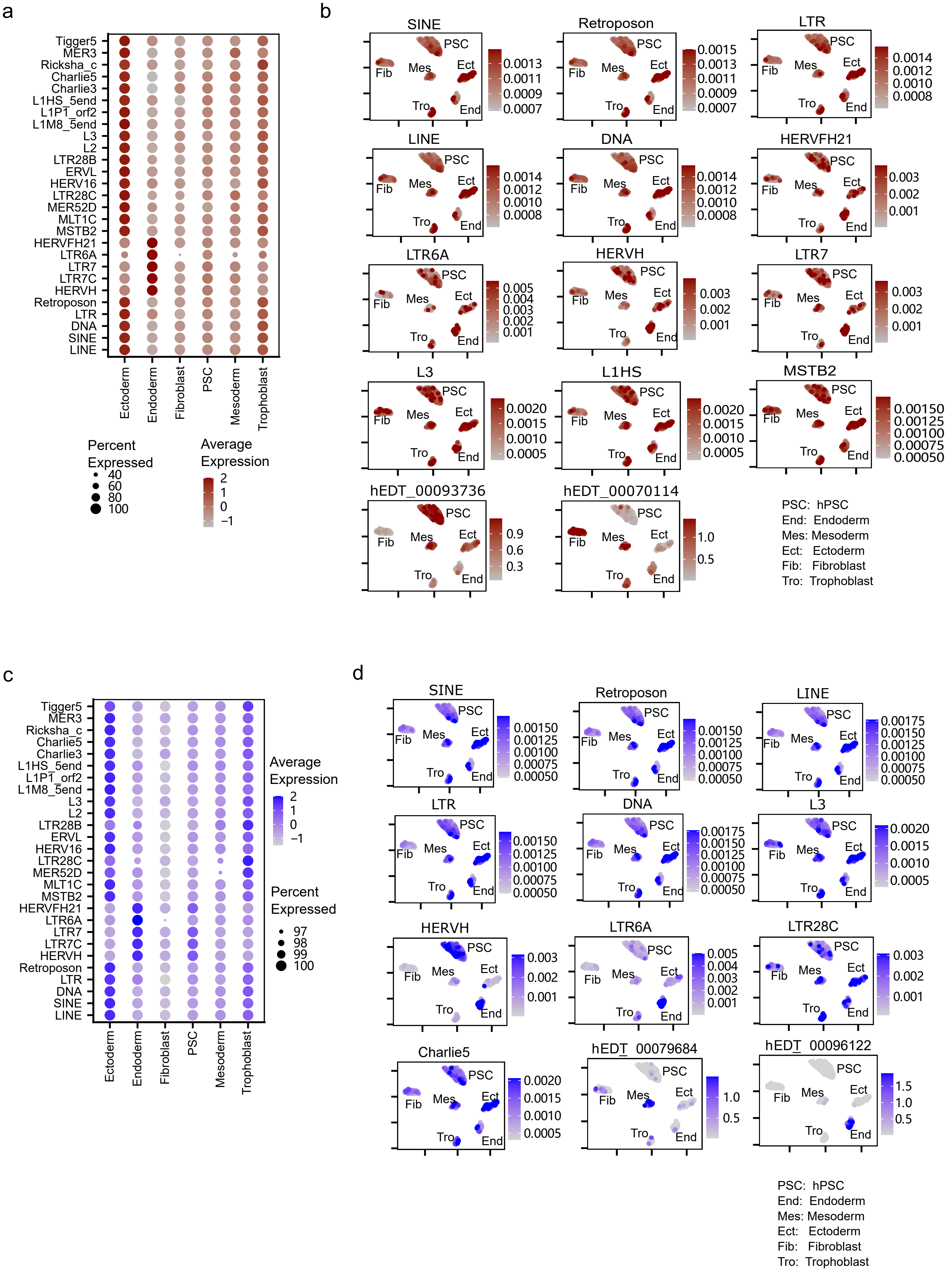


Extended Data Fig. 10

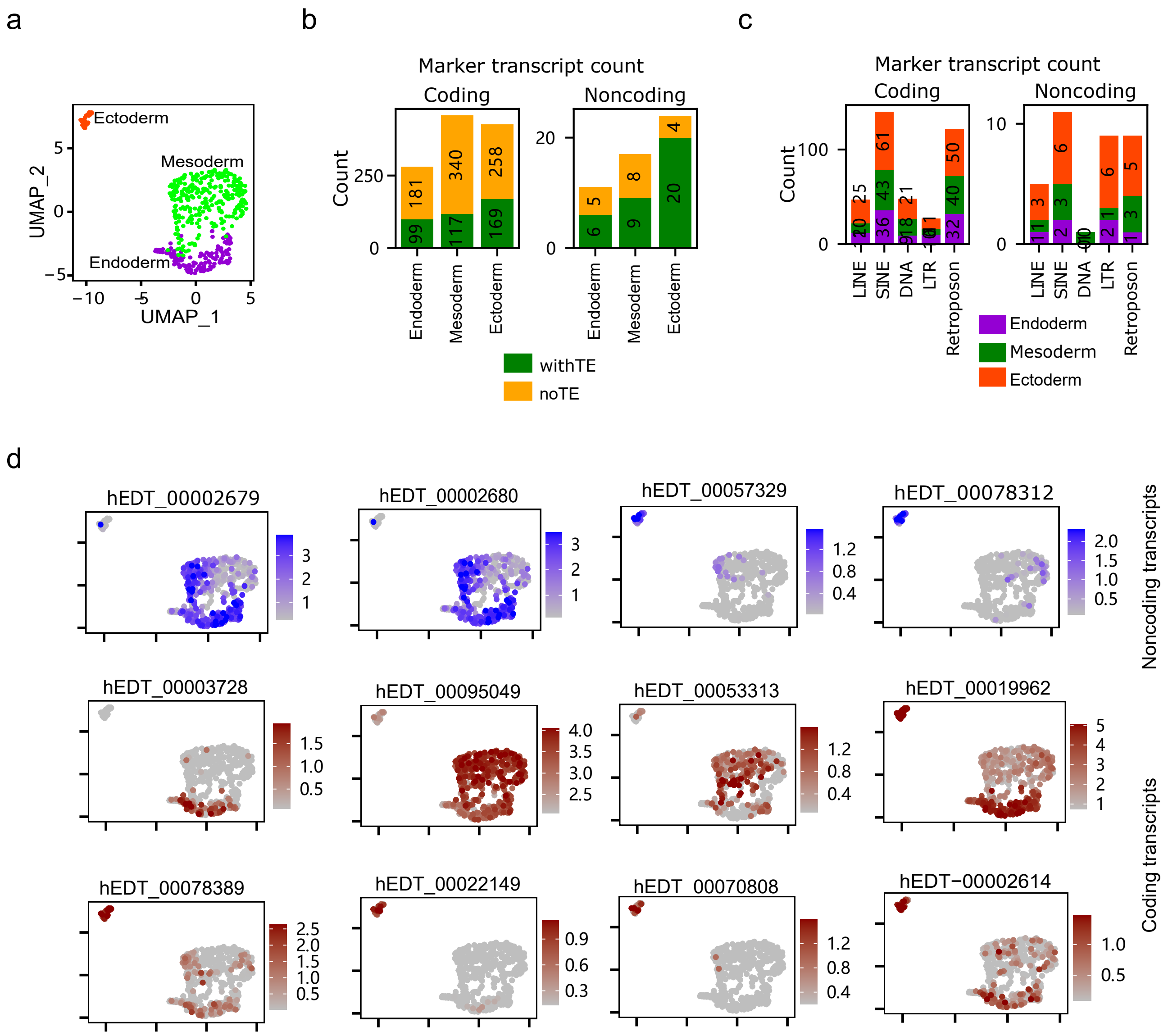


Extended Data Fig. 11

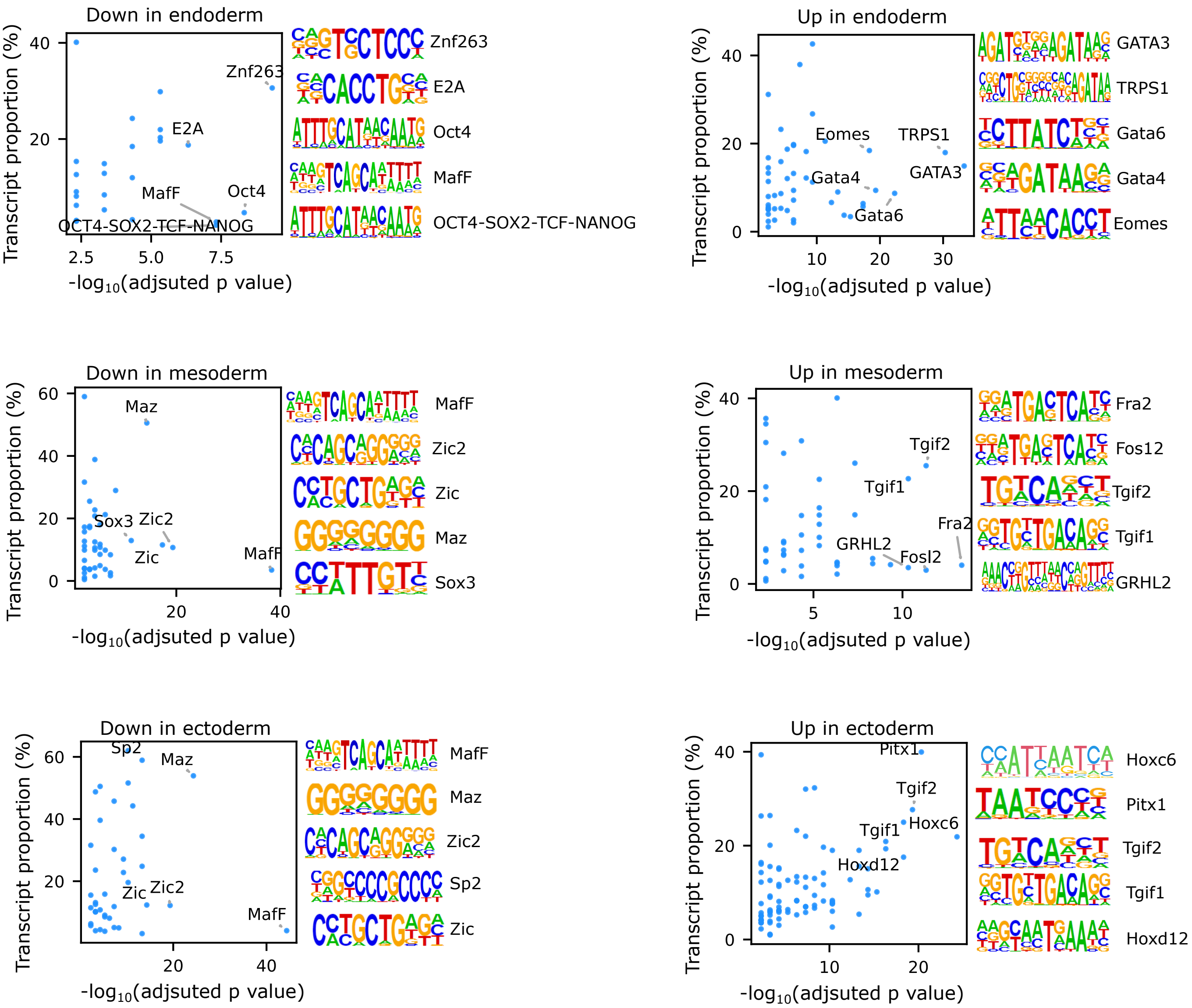


Extended Data Fig. 12

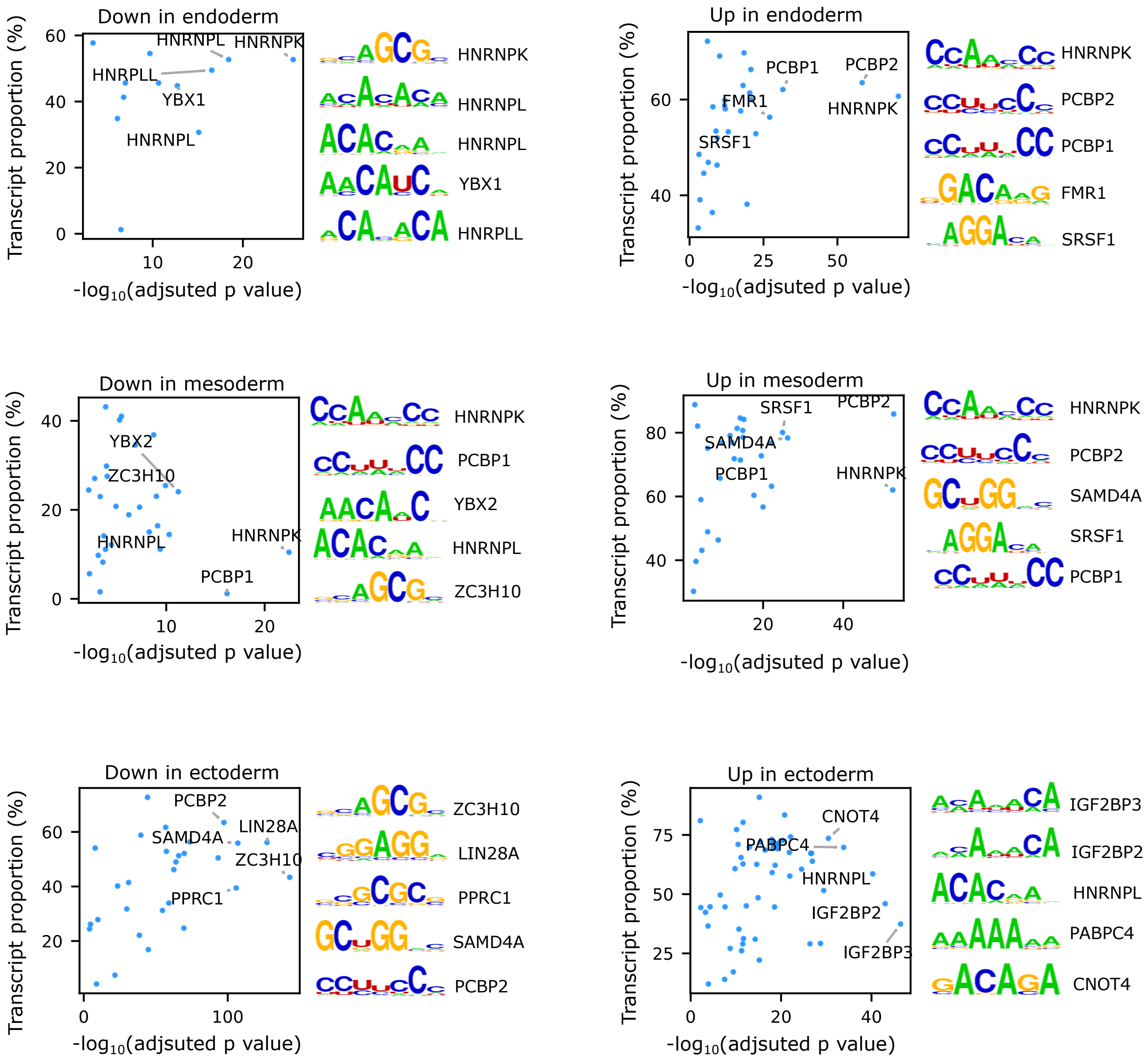


Extended Data Fig. 13

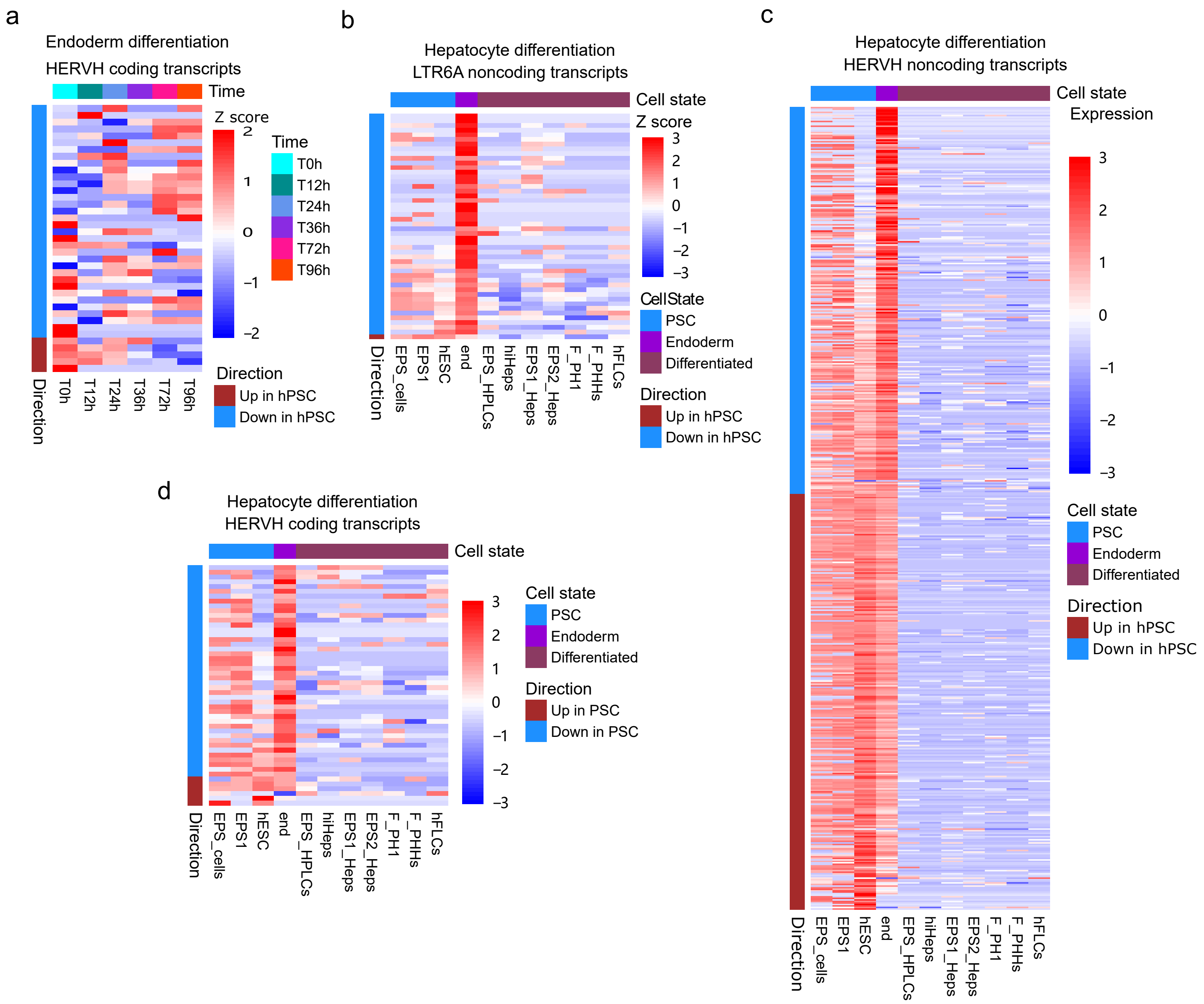


Extended Data Fig. 14

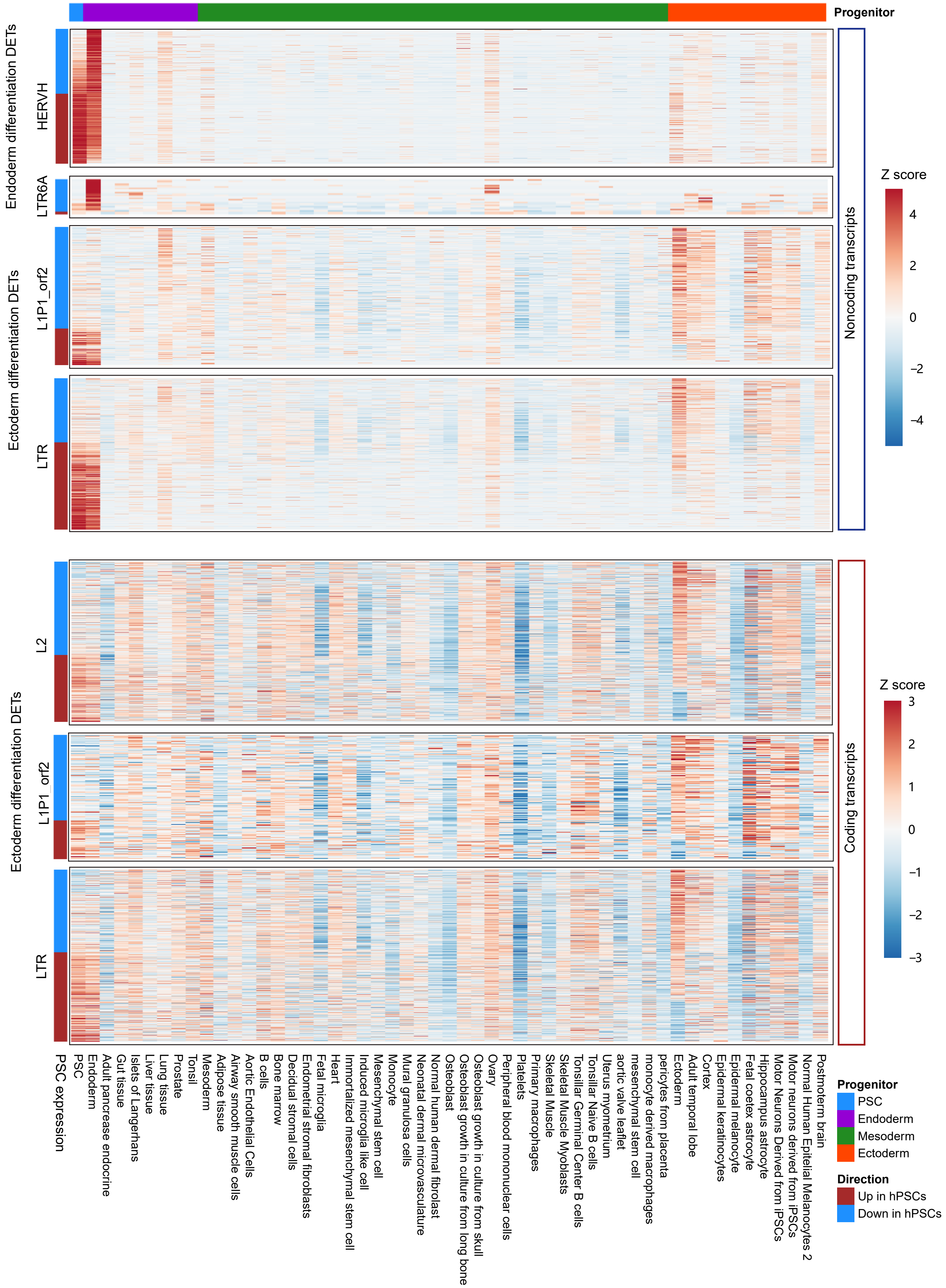


Extended Data Fig. 15

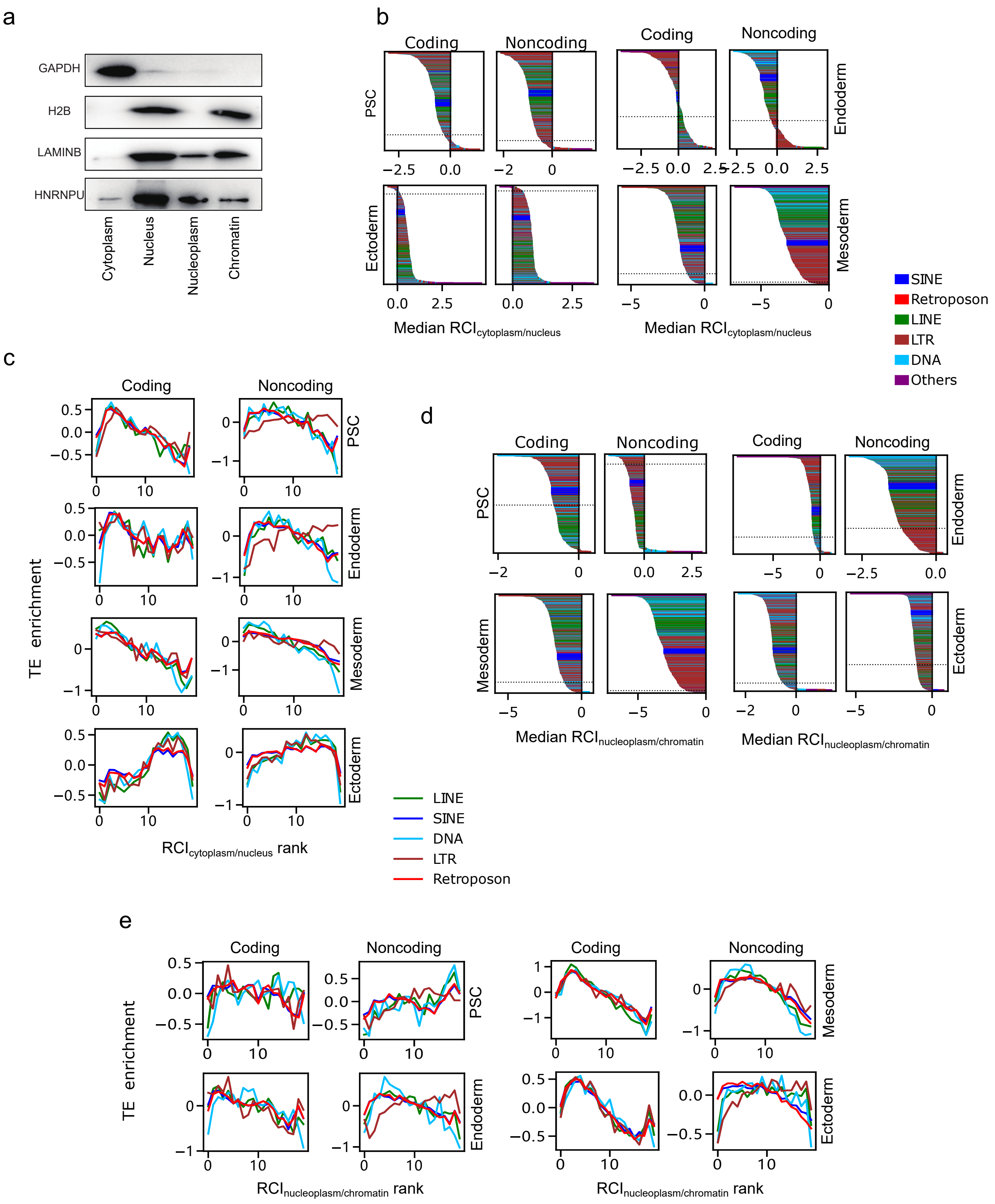


Extended Data Fig. 16

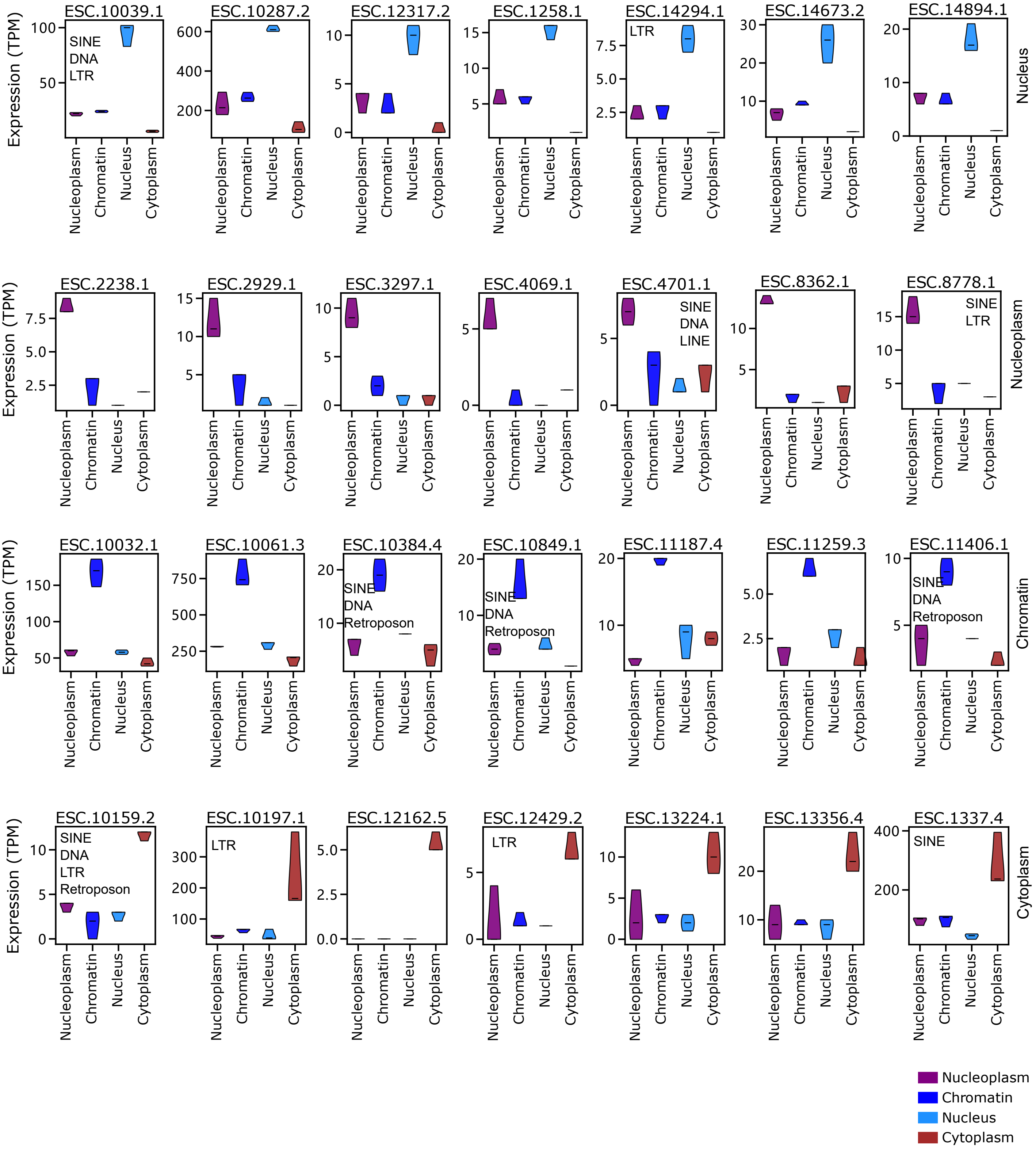


Extended Data Fig. 17

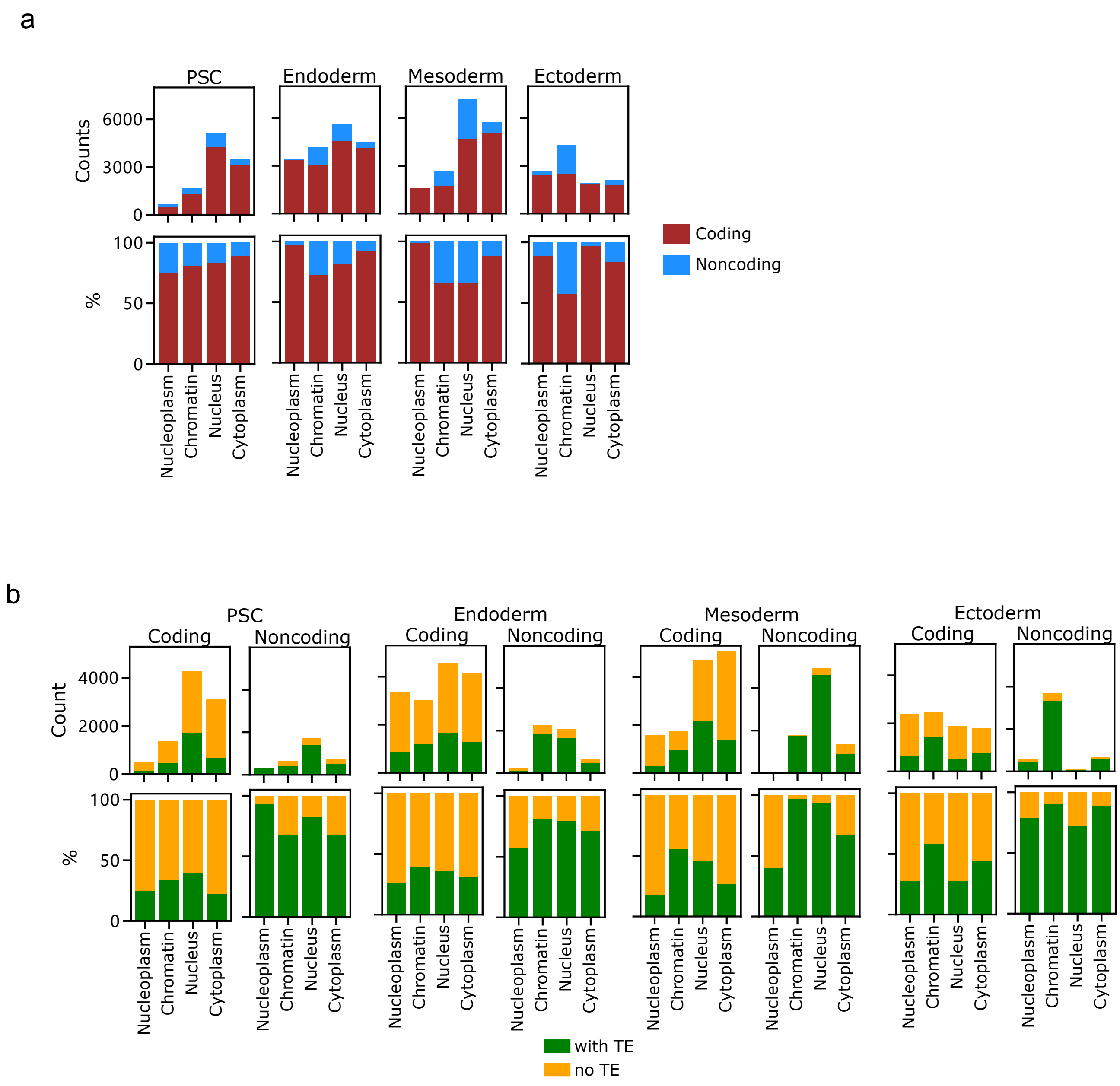


Extended Data Fig. 18

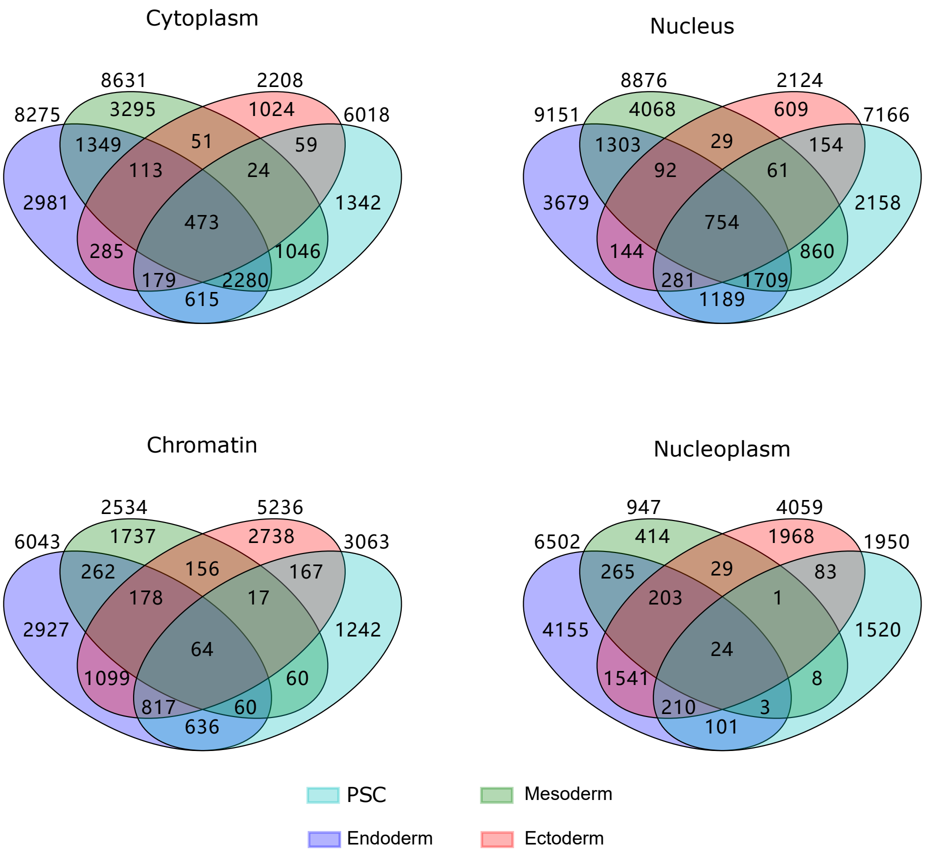


Extended Data Fig. 19

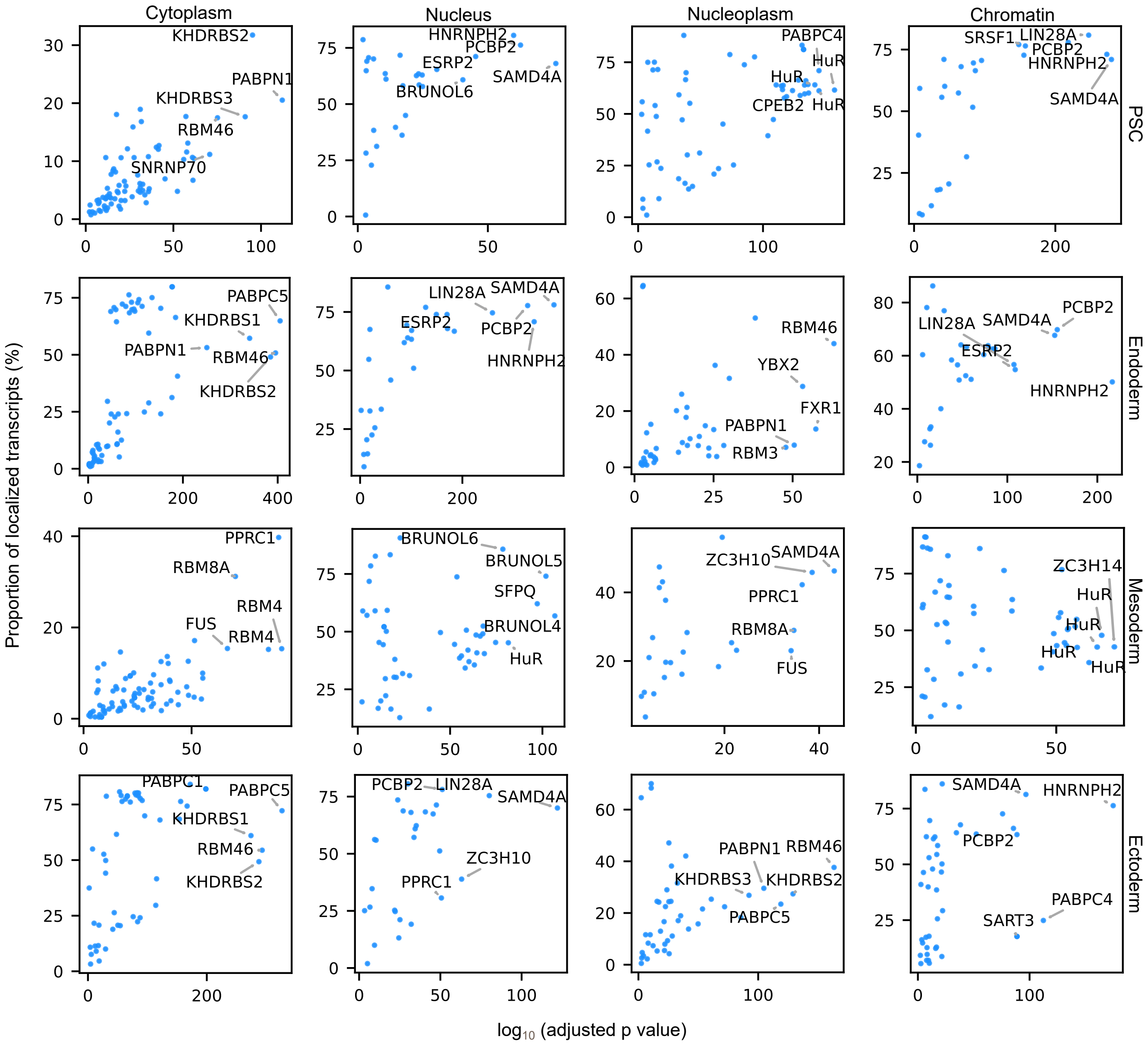


Extended Data Fig. 20

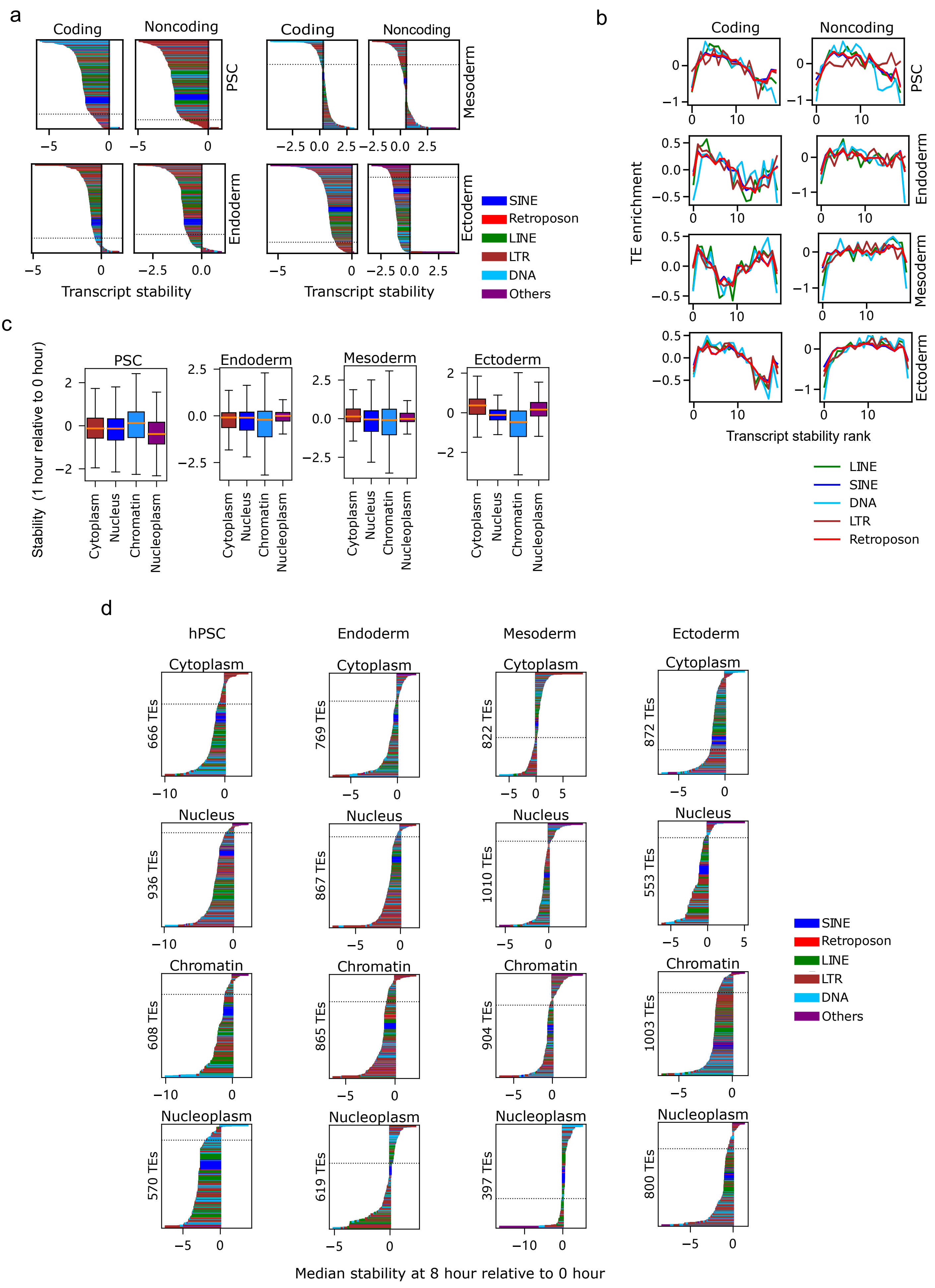


Extended Data Fig. 21
